# Conservation of dynamic characteristics of transcriptional regulatory elements in periodic biological processes

**DOI:** 10.1101/2020.10.12.328658

**Authors:** Francis C. Motta, Robert C. Moseley, Bree Cummins, Anastasia Deckard, Steven B. Haase

## Abstract

Cell and circadian cycles control a large fraction of cell and organismal physiology by regulating large periodic transcriptional programs that encompass anywhere from 15-80% of the genome. The gene-regulatory networks (GRNs) controlling these programs were largely identified by genetics and chromosome mapping approaches in model systems, yet it is unlikely that we have identified all of the core GRN components. Moreover, large periodic transcriptional programs controlling a variety of processes certainly exist in important non-model organisms where genetic approaches to identifying networks are expensive, time-consuming or intractable. Ideally, the core network components could be identified using data-driven approaches on the transcriptome dynamics data already available. Previous work used dynamic gene expression features to identify sets of genes with periodic behavior; our work goes further to distinguish genes by role: core versus their non-regulatory outputs. Here we present a quantitative approach that can identify nodes of GRNs controlling cell or circadian cycles across taxa. There are practical applications of the approach for network biologists, but our findings reveal something unexpected—that there are quantifiable and fundamental shared features of these unrelated GRNs controlling disparate periodic phenotypes.

**Author summary:** Circadian rhythms, cellular division, and the developmental cycles of a multitude of living creatures, including those responsible for infectious diseases, are among the many dynamic phenomena in the natural world that are known to be the eventual output of gene regulatory networks. Identifying the small number of specialized genes that control these dynamic behaviors is of fundamental importance to our understanding of life, and our treatment of disease, but is difficult because of the sheer size of the genomes. We show that the core genes in organisms separated by millions of years of evolution have remarkable similarities that can be used to identify them.

## Introduction

Periodic phenotypes span nearly the entire tree of life and include such fundamental processes as the cell-division cycle, circadian rhythms, and developmental cycles. Probing the genetic mechanisms that give rise to these dynamic activities is not only crucial to our fundamental understanding of life and its evolution, it will also add to the current collection of synthetic biology components and principles of design, and may reveal novel treatments for disease and infection. A vast body of experimental evidence, gathered over years of targeted experimentation (e.g. gene knock-outs) has uncovered the existence of endogenous circadian clocks: complex GRNs—comprised mostly of interacting transcription factors (TFs)—within cyanobacteria, fungi, plants and mammals [1–3]. Moreover, a GRN also appears to control the timing of cell-cycle events in budding yeast [4–8]. To understand the complex dynamic functions of these GRNs, experimentalists and computational scientists have developed a variety of approaches to infer the structure of GRNs. An essential first step is to identify, from among an expansive set of candidate genes, those *core* gene products controlling the dynamics of the associated program of gene expression. We conceptualize core nodes as interacting in a strongly connected subnetwork of mutual activation and repression. The core then drives the dynamics of “output” or “effector” nodes that do not feed back into the core but rather transmit the dynamic expression pattern to downstream target genes (Fig. 1).

**Fig 1.**
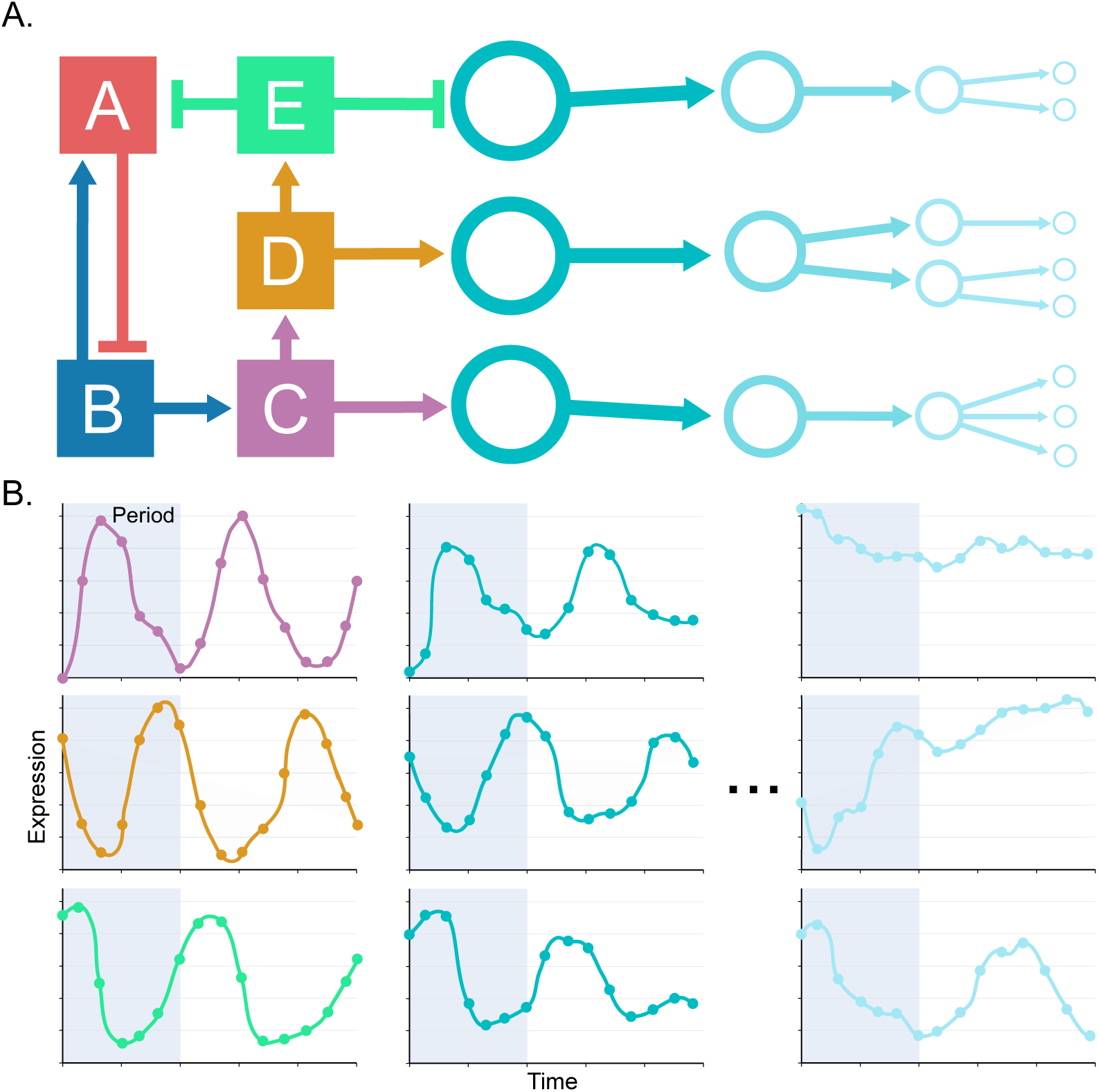
Conceptual Model of Core Regulatory Elements. (A) Conceptual model of a transcriptional regulatory network with core nodes (squares) operating in a strongly-connected subnetwork of mutual activation (arrows) and repression (short bars), together with outputs of the core (circles). Output nodes transmit the transcriptional signal that is generated by the core, but which diminishes as it moves away from core nodes. (B) Illustrations of transcript abundance profiles exhibited by the core and its output nodes, with core nodes having oscillations that have a precise match to a specified period (shaded region) and large variations in expression.

Identifying core nodes is especially daunting for organisms where genetic experiments are largely intractable. Moreover, functional redundancy, and complex GRN mechanisms, such as accessory feedback loops, can complicate the discovery of core nodes. Here we identify distinguishing characteristics of the dynamics of gene expression that are conserved across organisms that are separated by hundreds of millions of years of evolution, in vastly different biological processes, and across data-collection modalities. We discover that a combination of dynamic features provides a rank ordering of all genes such that core nodes are generally highly-ranked, even among the many genes which exhibit these features. Moreover, we find that, in general, a combination of dynamic features more accurately distinguishes core transcriptional regulators than individual features on their own. Our findings support the use of quantified dynamic characteristics of gene expression to identify core regulatory elements and show that there are common features in the dynamic gene expression of core regulatory variables that drive very different biological processes.

## Materials and Methods

### Dynamic Curve Features

We utilize quantified measures of 1) periodicity at a specified period, and 2) regulatory strength associated to each time series transcript abundance profile across a transcriptome. The metrics used in this study are summarized in Table 1. Detailed descriptions of the algorithms used to compute these metrics are available in the Supplementary Materials.

**Table 1.** Quantitative metrics of periodicity and regulation strength used in this study to rank genes. *Refer to the Supplementary Materials for equation definitions.

| Name | Function | Type | Description |
| --- | --- | --- | --- |
| DL Per Score | Per(G) | periodicity | A measure of abundance profile periodicity as defined by Eq. 3*. |
| DL Per $p$-val | $p_{per}(G)$ | periodicity | An empirical $p$-value measuring the probability that a random abundance profile will exhibit a DL Per Score larger than the actual gene’s expression pattern. |
| JTK Per $p$-val | $p_{jtk}(G)$ | periodicity | An analytic $p$-value introduced in [9] measuring the correlation in the discrete up-down patterns of expression between a gene and a sinusoidal template. |
| DL Reg Score | Reg(G) | regulation | A measure of the variability of transcript abundance about its mean expression level as defined by Eq. 2* |
| DL Reg $p$-val | $p_{reg}(G)$ | regulation | An empirical $p$-value measuring the probability that a random abundance profile will exhibit a DL Reg Score larger than the actual gene’s. |
| PerReg |  | combined | The product of DL Per and DL Reg Scores. |
| DL |  | combined | The original periodicity measure introduced in [10] and defined according to Eq. 1*. |
| DL×JTK | | combined | A modified version of the original periodicity measure introduced by [10], defined according to Eq. 1* with $p_{per}(G)$ replaced by $p_{jtk}(G)$. |

### Performance of Gene Ranking Metrics

The problem of identifying the core regulatory elements within an organism’s genome is fundamentally a question of binary classification of gene function: is a gene core or not? In practice, this decision task amounts to ranking all genes by some quantitative metric or “score” in the hope that the ranking is enriched with core genes, so as to reduce the expected effort required to gather additional experimental evidence of gene function through, for example, knock-out experiments.

To assess the capacity of each ranking metric given in Table 1 to distinguish core from non-core genes, we compute the precision-recall (PR) curves of the gene rankings. PR curves track the precision (the fraction of true core genes among all genes ranked above some score threshold) across all levels of recall (the fraction of true core genes appearing above the chosen threshold). From each PR curve we compute the average precision (AP), which summarizes with a single number a ranking’s performance across all recall levels. See the Supplementary Materials for a more complete description of PR curves, precision, recall and AP.

For us, a perfect ranking of genes is one in which all core genes are ranked higher than all non-core genes. In this way, an experimentalist prioritizing hypotheses using the gene ranking would encounter all core genes before testing any non-core. The AP of a perfect ranking will be 1. At the other extreme is an uninformative ranking which assigns scores to genes at random. The average precision achieved for a random classifier is *C/N* [11], where *C* is the number of core genes and *N* is the number of all genes. Moreover, the expected PR curve for such an algorithm is a horizontal line at precision level *C/N* across all recall levels, as seen in Figs. S1–S6. Thus, performance of each classifier, as measured by its PR curve and its AP, should be compared against the (non-universal) baseline performance of a random classifier. In other words, precision-recall points above the baseline reflect the skill of a metric, over the random classifier, to rank genes in a way which enriches the top of a list with core genes.

### Gene Expression Datasets

#### Data Processing

The normalized transcriptomic datasets used in this analysis were taken from the references presented in Table 2. Before deriving dynamic features, transcript abundances were preprocessed to remove unreliable data. For the *M. musculus* and *S. cerevisiae* RNAseq datasets, genes whose normalized transcript levels were less than 1 FPKM for more than half of their time points were dropped from the dataset and not considered in any part of this analysis.

**Table 2.**
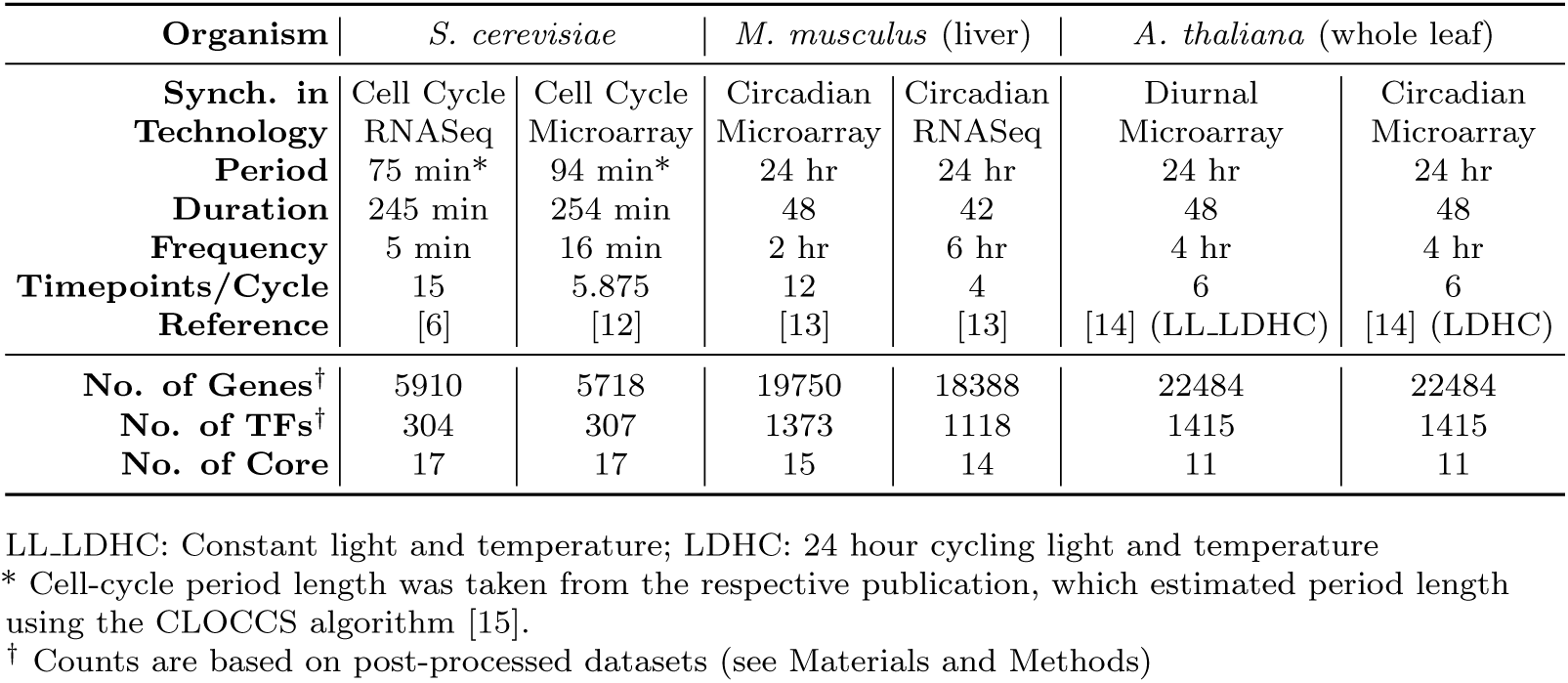
Time series transcript abundance datasets used in this study.

| Organism | *S. cerevisiae* |  | *M. musculus* (liver) |  | *A. thaliana* (whole leaf) |  |
| --- | --- | --- | --- | --- | --- | --- |
| **Synch. in** | Cell Cycle | Cell Cycle | Circadian | Circadian | Diurnal | Circadian |
| **Technology** | RNASeq | Microarray | Microarray | RNASeq | Microarray | Microarray |
| **Period** | 75 min* | 94 min* | 24 hr | 24 hr | 24 hr | 24 hr |
| **Duration** | 245 min | 254 min | 48 | 42 | 48 | 48 |
| **Frequency** | 5 min | 16 min | 2 hr | 6 hr | 4 hr | 4 hr |
| **Timepoints/Cycle** | 15 | 5.875 | 12 | 4 | 6 | 6 |
| **Reference** | [6] | [12] | [13] | [13] | [14] (LL-LDHC) | [14] (LDHC) |
| **No. of Genes<sup>†</sup>** | 5910 | 5718 | 19750 | 18388 | 22484 | 22484 |
| **No. of TFs<sup>†</sup>** | 304 | 307 | 1373 | 1118 | 1415 | 1415 |
| **No. of Core** | 17 | 17 | 15 | 14 | 11 | 11 |

LL-LDHC: Constant light and temperature; LDHC: 24 hour cycling light and temperature
\* Cell-cycle period length was taken from the respective publication, which estimated period length using the CLOCCS algorithm [15].
<sup>†</sup> Counts are based on post-processed datasets (see Materials and Methods)

Authors of [6] produced the *S. cerevisiae* microarray dataset from *S. cerevisiae* cells that were synchronized via centrifugal elutriation. It is known that elutriation impacts the transcription of many genes and that a brief recovery period is needed after elutriation. The resulting transcript abundance dynamics early in the time series, which are not related to cell-cycle transcript abundance dynamics, can impact periodicity analyses [15]. Therefore, prior to any analysis, [6] eliminated data determined to be associated with the elutriation recovery period. We adopted the same method of eliminating the first two time points from the *S. cerevisiae* microarray dataset.

In the *S. cerevisiae* mircoarray dataset and both *A. thaliana* datasets, some genes were associated with multiple probes, causing some genes to have more than one transcript abundance profile. The *A. thaliana* core gene, *RVE8*, was one such gene. Having two transcript abundance profiles for *RVE8* resulted in inaccurate performance metrics. To remedy this issue, we applied a filtering step to the *S. cerevisiae* mircoarray dataset and both *A. thaliana* datasets after quantifying dynamic features using the methods in Table 1. For genes with multiple abundance profiles, we kept the profile with the highest average abundance, resulting in the elimination of 96 and 326 profiles from the *S. cerevisiae* mircoarray dataset and both *A. thaliana* datasets, respectively. All time series data can be found in Dataset S1.

#### Curation of Core Regulatory Elements

In order to evaluate the ability of each method given in Table 1 to identify core TFs driving a periodic program of gene expression, we consider data derived from well-studied organisms for which there is significant experimental evidence of gene function. Core cell-cycle TFs in yeast are described as genes functioning in an autoregulatory transcriptional network that robustly maintains a large program of periodic gene expression [4–6, 8]. A list of yeast core cell-cycle TFs based on this definition was compiled in [16] for evaluating the transcriptonal oscillator underlying the yeast cell cycle. Therefore, the core TF list defined in [16] was used in this study as the ground truth for *S. cerevisiae* (Dataset S2). Similarly, core circadian clock TFs are described as genes functioning in an autoregulatory transcriptional feedback loop, maintaining circadian-like transcript abundances under constant light or dark conditions and are necessary components for generation and regulation of circadian rhythms [1, 17, 18]. The literature evidence supporting our labeling of plant and mammalian genes as core are listed in Dataset S2. Although the core networks are known to include non-TF regulatory elements that control functional activity, such as kinases and ubiquitin ligases [1, 18, 19], we limit our definition of core to TFs since these are more reliably annotated in the genomes we consider. This ensures our conclusions are conservative by not unfairly inflating the core list with known core post-transcriptional modifiers while not simultaneously including all non-core members of these gene categories.

#### Curation of Transcription Factors

In this study, we define a TF as a gene that has the ability for sequence-specific DNA binding alone or in a complex and is capable of activating and/or repressing gene expression. This definition excludes genes that are also known to affect gene expression, such as chromatin-related genes like chromatin remodeling factors, histone demethylases, and histone acetyltransferases. To ensure the lists of TFs are consistent across strains, we used curated TF databases that use the given TF definition. In particular, TFs used in this study (Dataset S3) were retrieved from Animal TF Database 3.0 [20], Plant TF Database 4.0 [21], and YEASTRACT [22] for *M. musculus, A. thaliana*, and *S. cerevisiae*, respectively. Each species list of TFs was inspected for presence of the respective species core regulatory elements. Upon inspection of the *A. thaliana* TF list, it was discovered that the core regulatory elements from the pseudo-response regulator (PRR) family were not present. Therefore, we added *PRR5, PRR7, PRR9*, and *PRR1* (*TOC1*) to *A. thaliana* list of TFs, which are known as core regulatory elements of the plant circadian clock [23–25].

## Results and Discussion

Understanding the function of GRNs requires a specification of the control variables and their interactions. Accurate inferences have generally required substantial genetic perturbation and physical localization studies and thus has been confined to experimentally tractable model systems. However, previous work has indicated that interactions between GRN nodes can be inferred directly from transcriptome dynamics data [16]. Here we investigated whether the list of core nodes could also be identified from time series transcriptomics. We determined that quantifiable features from time-series gene expression measurements can be used to identify experimentally-inferred core nodes from model systems across taxa (yeast cell cycle, mouse circadian cycle, plant circadian cycle).

We consider two quantifiable characteristics of dynamic transcript abundance profiles, measured in multiple ways, and assess the capacity of each to differentiate core from non-core regulatory elements. Because the dynamic phenotypes of interest are rhythmic, e.g. sleep-wake cycles, cell division, etc., it is natural to ask to what extent, relative to all genes, will the core elements driving these processes be endowed with periodicity that matches the observed cycling at the level of their transcript abundance? Moreover, since the core elements are by definition those TFs governing the dynamics of gene expression, to what extent will the strength of the regulatory signal be reflected in the dynamics of transcript abundance?

### Dynamic transcript abundance features identify regulatory elements in core networks

We first examined the list of dynamic features, used both individually and in various combinations (see Table 1) to distinguish core TFs from among all TFs. To provide a unified measure of performance across datasets we considered the AP of each metric’s ranking of transcripts. When restricting to TFs, using both periodicity and regulation strength features together yields significantly higher AP scores than the baseline for each of the six datasets examined (Fig. 2A). Even using just one of the two types of dynamic features, we see remarkable improvement over baseline, although generally lower AP scores, than the combined metrics, across all six datasets (Fig. 2B). These results are significant since the datasets considered in this study represent organisms from three different kingdoms, undergoing two ostensibly mechanistically distinct periodic dynamic processes. The complete set of metrics scoring all genes in all datasets are available in Dataset S4 and the complete precision-recall curves for all datasets and all metrics are available in Figs. S1–S6.

**Fig 2.**
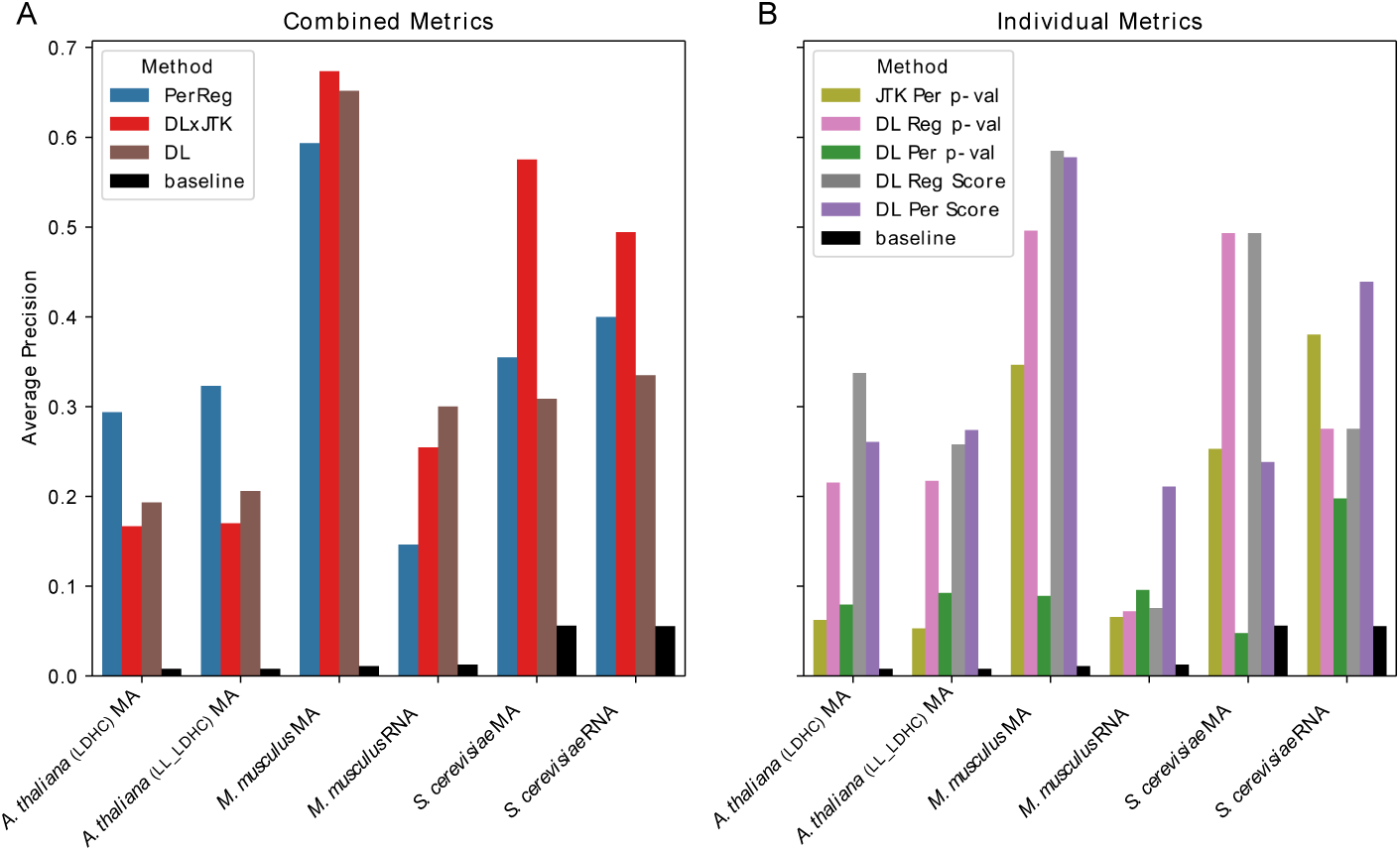
Identifying Core Genes Among Transcription Factors. Average precision of classifiers identifying core from non-core TFs among all TFs by combined metrics (A) and individual metrics (B) (Table 1) as well as the baseline average precision of a random classifier, for each dataset (Table 2).

From the viewpoint of an experimentalist interested in understanding the entirety of a core network, it is encouraging to observe the enrichment of the top 25 TFs with core genes. Among the top 25 TFs ranked by the measure DL × JTK, 13 (12) of the possible 17 *S. cerevisiae* core genes are identified using the microarray (RNASeq) data. Similarly, 10 (4) core *M. musculus* genes from the possible list of 15 (14) core genes, are among the top 25 transcription factors as ranked by DL × JTK using microarray (RNASeq) data. Finally, *A. thaliana* LDHC and LL LDHC datasets contain 4 and 5 core genes, respectively, from among the 11 possible core, in the top 25. Perhaps even more amazingly, 9 of the top 10 *M. musculus* TFs and 6 of the top 10 *S. cerevisiae* TFs are core when the high temporal resolution microarray datasets are ranked using DL × JTK. These results are given in Table 3.

**Table 3.** Top 25 transcription factors ranked by DL×JTK metric.

| Rank | *S. cerevisiae* |  | *M. Musculus* |  | *A. thaliana* |  |
| --- | --- | --- | --- | --- | --- | --- |
|  | MA | RNA | MA | RNA | LDHC | LL_LDHC |
| 1 | **SWI5\*** | **TOS4\*** | **ARNTL\*** | **DBP\*** | *COL1* | *STH* |
| 2 | **YOX1\*** | *HST4* | **DBP\*** | **NPAS2\*** | *HB-12* | *AT1G26790* |
| 3 | *HST3* | *HST3* | **NPAS2\*** | *CDX4* | *TGA3* | **CCA1\*** |
| 4 | *ASF1* | **SWI5\*** | **NR1D1\*** | **ARNTL\*** | *RVE1* | *BBX18* |
| 5 | **ACE2\*** | **YOX1\*** | **NR1D2\*** | *EGR1* | *MYBL2* | *COL1* |
| 6 | *RTT107* | *RTT107* | **BHLHE41\*** | *GM14401* | **LHY\*** | *CDF1* |
| 7 | **STB1\*** | *WTM2* | **CLOCK\*** | *GM14305* | *CO* | *COL2* |
| 8 | **HCM1\*** | **ASH1\*** | **NFIL3\*** | *POU4F1* | *PIL6* | *CDF3* |
| 9 | *RME1* | **FKH1\*** | *RFXANK* | *EN2* | *AT2G28200* | *AT2G28200* |
| 10 | **FKH1\*** | *ASF1* | **RORC\*** | *DMRTA2* | *COL2* | *RVE1* |
| 11 | **PLM2\*** | **ACE2\*** | **TEF\*** | *LHX1* | **CCA1\*** | **LHY\*** |
| 12 | **SWI4\*** | *POG1* | *CREM* | *GM20422* | **PRR7\*** | *COL5* |
| 13 | **NDD1\*** | **SWI4\*** | *EGR1* | *GM14444* | *HYH* | *PIF4* |
| 14 | **ASH1\*** | *RME1* | *PPARD* | *OVOL2* | *BBX18* | *PIL6* |
| 15 | **YHP1\*** | **PLM2\*** | *ZBTB21* | *GM4969* | **RVE8\*** | *BBX16* |
| 16 | **TOS4\*** | *RLF2* | *NFIC* | *HOXC4* | *PRE1* | **LUX\*** |
| 17 | *EDS1* | **NDD1\*** | *AHCTF1* | *FOXO6* | *BZS1* | **PRR7\*** |
| 18 | *RIF1* | **HCM1\*** | *ATF5* | *MESP1* | *EPR1* | *CDF2* |
| 19 | *SIP4* | *GAT1* | *LITAF* | *AI854703* | *CDF3* | *LZF1* |
| 20 | **FHL1\*** | *TEC1* | *KLF10* | **NR1D1\*** | *RVE2* | *HB-12* |
| 21 | *NUT1* | **STB1\*** | *KLF13* | *BNC2* | *AT1G26790* | **RVE8\*** |
| 22 | *ASG1* | **YHP1\*** | *ESR1* | *NPAS3* | *BBX16* | *ATCTH* |
| 23 | *TBF1* | *RPI1* | *STAT5B* | *2210418O10RIK* | *COL9* | *MYBL2* |
| 24 | *SNF5* | *MTH1* | *SREBF1* | *HOXC6* | *LZF1* | *ARF11* |
| 25 | *WTM2* | *RIF1* | *MAFB* | *TBX1* | *ARF10* | *RL6* |
| Recall | 76.5% | 70.6% | 66.7% | 28.6% | 36.4% | 45.5% |

LL\_LDHC: Constant light and temperature; LDHC: 24 hour cycling light and temperature; MA: Microarray; RNA: RNAseq
\* Core transcription factors in Dataset S2

We emphasize the skill of dynamic gene expression features to identify core TFs in Fig. 3, which gives the distribution of core TF DL×JTK ranks among all TFs for *S. cerevisiae* (see also Tab. S1) and heatmaps of microarray gene expression grouped by DLxJTK rankings. The top 25 genes are clearly seen to robustly oscillate at approximately the specified period (94 min) and among these are 13 of the 17 core genes.

**Fig 3.**
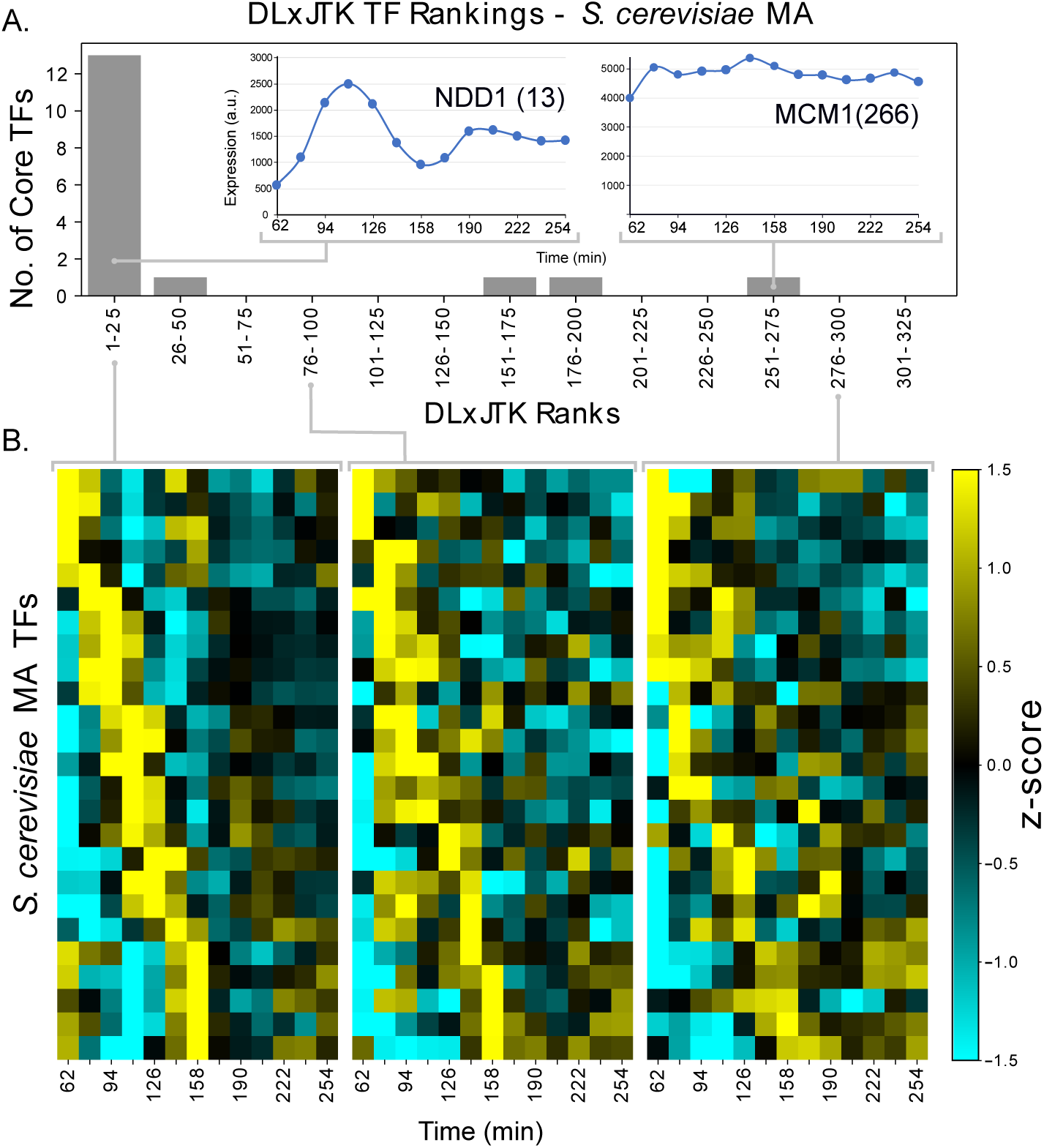
Transcript abundance dynamics across DL × JTK rankings of transcription factors. (A) Distribution of DL × JTK ranks of core *S. cerevisiae* TFs among all TFs and time series expression of two core TFs: NDD1, which is highly ranked (rank 13), and MCM1, which is not highly ranked (rank 266). NDD1 and MCM1 act in a complex to regulate downstream targets. (B) Heatmaps of standardized gene expression profiles of the genes ranked (left) 1-25, (middle) 76-100, and (right) 276-300 by DL×JTK. Within each subpanel, genes are ranked by peak expression.

The recall of core genes by DL × JTK among the top 25 TFs is as much as 76.5% of the core yeast cell-cycle transcriptional regulatory network, up to 66.67% for the mouse circadian clock with well-sampled data, and 45.45% for the core plant circadian network under circadian conditions. Meaning, by using only the dynamics of transcript abundance and a list of TFs, an experimentalist would identify three-quarters of the known core cell-cycle TFs in yeast, two-thirds of the core circadian TFs in mice, and almost half of the core circadian TFs in plants from among the top 25 TFs when ranked using a combined measure of periodicity and regulation strength. Other combined measures perform skillfully when examining the top 25 ranked TFs, although not as consistently well across all the datasets as DL × JTK (Tables S2 and S3).

The ability of dynamic characteristics to identify core TFs from among all TFs may depend on the data collection modality and will certainly depend on the number of time points per cycle collected. This is made apparent by comparing the *S. cerevisiae* RNASeq and microarray datasets and, separately, *M. musculus* RNASeq and microarray datasets. We expect that the reduced DL × JTK classifier performance is largely due to the sensitivity of the JTK algorithm to the number of timepoints per cycle [26], although we cannot conclusively rule out the impact of the data type.

At the same time, quantitative measures of rhythmicity in transcript abundance and strength of regulation both independently improve the skill of a classifier above random. Thus, the functional regulatory elements driving very different biological processes exhibit common characteristics in the dynamics of their transcript expression.

### Dynamic transcript abundance characteristics remain adept at identifying core regulatory elements, even in the absence of prior knowledge of transcription factors

The organisms chosen for this study are model organisms in mammalian, plant, and fungi research which have been extensively studied. Thus, for these organisms, there are reliable annotations of gene function and comprehensive lists of TFs. If studying a non-model organism, evidence of gene function may be much weaker, for example relying on sequence-based inferences. We ask, to what extent do the dynamic characteristics of transcript abundance that distinguish core TFs from non-core TFs continue to distinguish core from all genes? In this way, we assess the capacity for gene expression dynamics to reduce hypothesis space in the absence of any prior biological knowledge. Note, this is an extremely lofty goal given the minuscule fraction of these genomes occupied by core transcriptional regulator elements.

For each dataset in Table 2 we ranked all transcript abundance profiles using the methods in Table 1. We have chosen to be very conservative in our labelling of core genes: only 17 out of nearly 6000 transcripts in *S. cerevisiae*, 14 out of close to 20,000 genes in *M. Musculus*, and 11 of over 22,000 genes in *A. thaliana*. As expected, AP scores are greatly reduced across all datasets. However, the APs remain significantly above baseline in most cases (Fig. 4). Examining the top 25 genes ranked by the measure DL×JTK, at least one core TF remained in the top 25 for all datasets, except the *A. thaliana* LDHC microarray dataset (Dataset S4). Remarkably, six of the 15 core mouse circadian TFs (recall of 40%) are identified among the top 25 genes ranked by DL×JTK in the *M. Musculus* liver microarray dataset.

**Fig 4.**
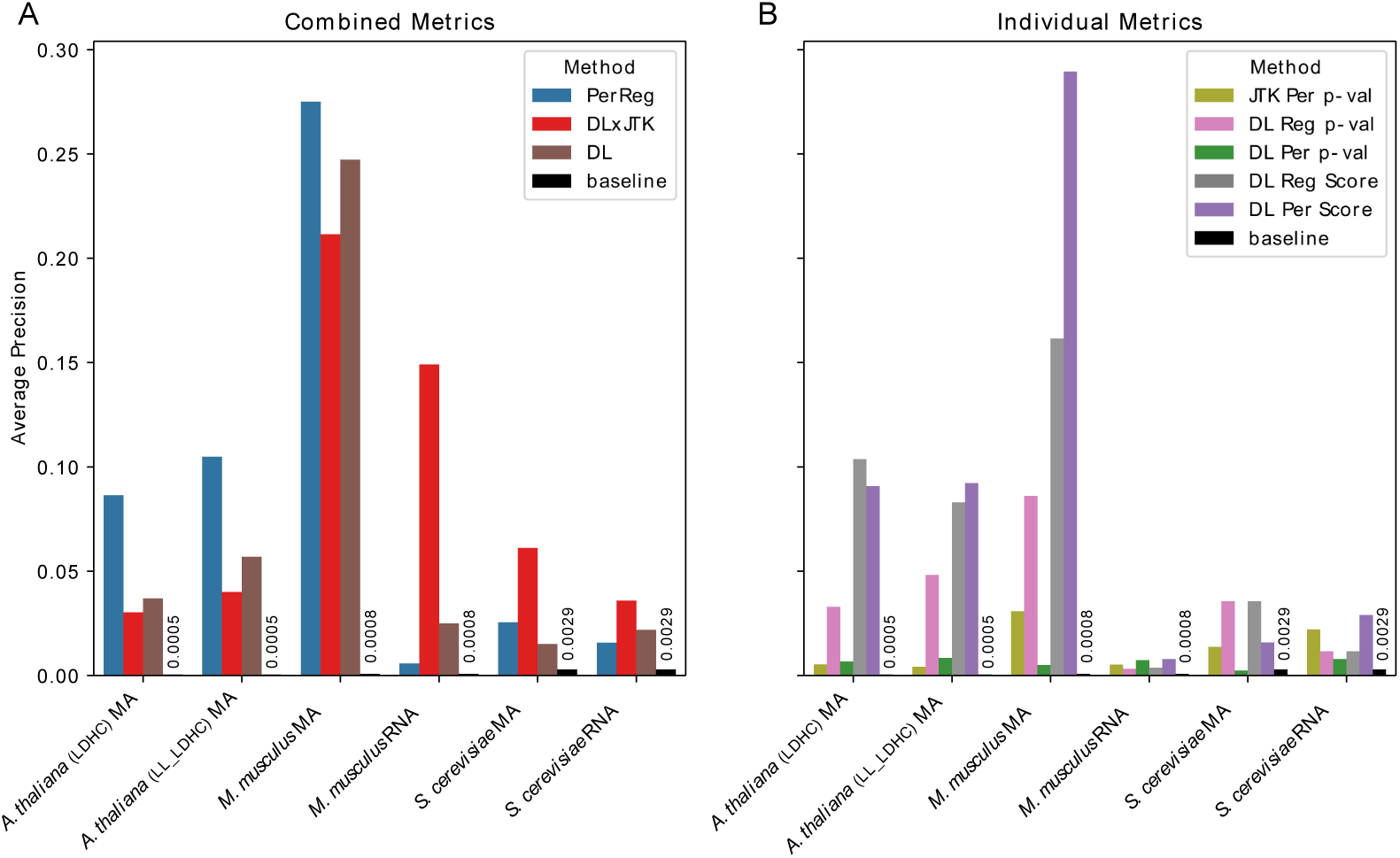
Identifying Core Genes Among All Genes. Average precision of classifiers identifying core from non-core TFs among all genes by combined metrics and individual metrics (Table 1) as well as the baseline average precision of a random classifier, for each dataset (Table 2).

### The dynamic transcript abundance characteristics of core regulatory elements are not overrepresented among transcription factors

It is certainly possible that the dynamic features under investigation are characteristic of TFs themselves, and thus our filtering on TFs causes us to already select for these features. To investigate the possibility that the dynamic metrics in this study are overrepresented in TFs and not just core transcriptional regulatory elements, we assessed the ability of the dynamic characteristics of transcript abundance to identify TFs from among all transcripts. In line with our hypothesis, all methods listed in Table 1 performed poorly as each method’s AP dropped to near or below the AP baseline (Fig. 5). Said another way, TFs within these organisms are effectively randomly distributed in the rankings of all genes by periodicity and variability of transcript abundance. The inability of the methods to identity TFs in each dataset demonstrates that these dynamic features are not characteristic of TFs in general, although they are indicative of core regulatory elements.

**Fig 5.**
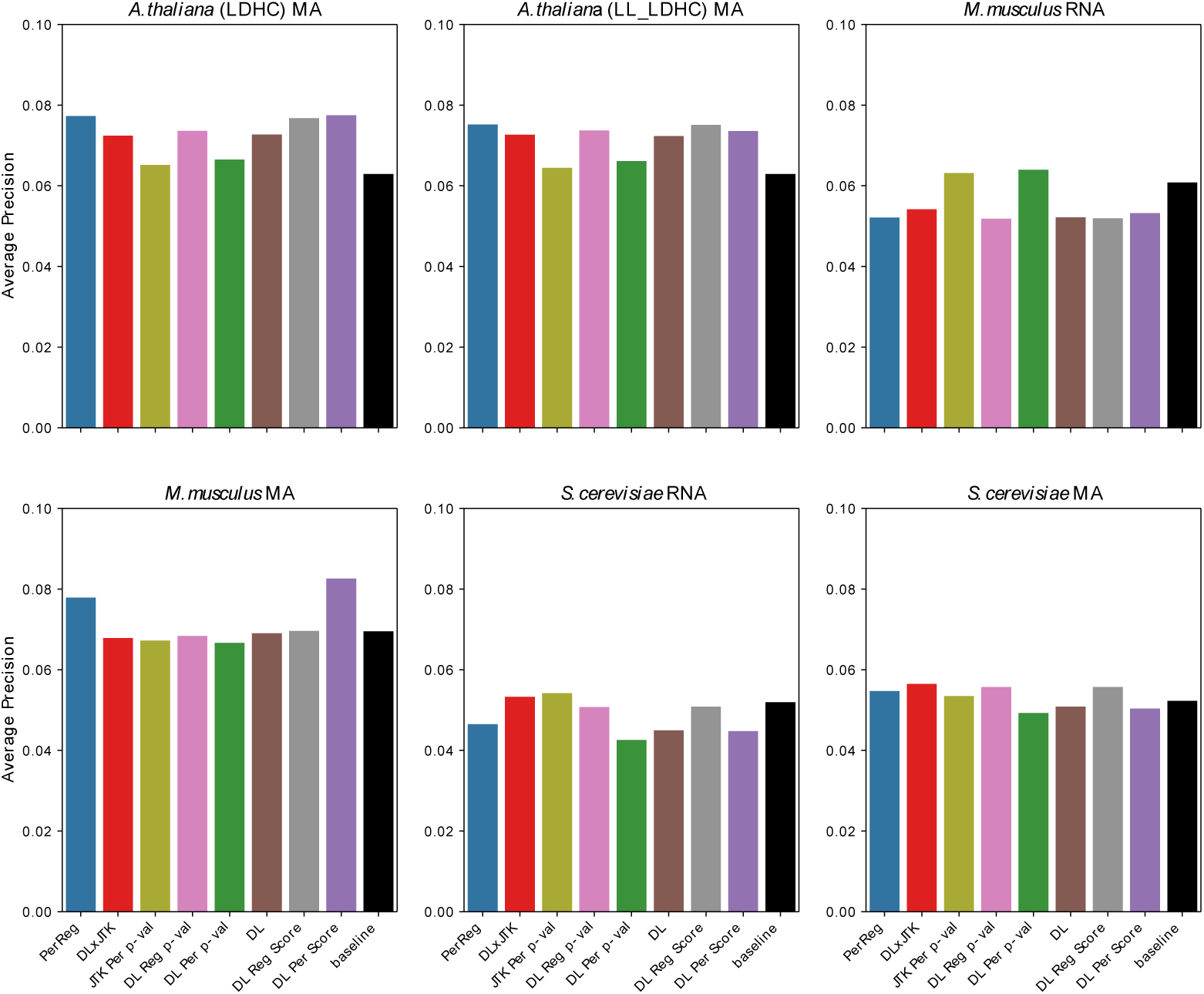
Identifying Transcription Factors Among All Genes. Average precision of classifiers identifying TFs from non-TFs among all genes by combined metrics and individual metrics (Table 1) as well as the baseline average precision of a random classifier, for each dataset (Table 2).

### Statistical significance measures are not required to skillfully rank core genes

A major concern with the DL methods for determining significance is that they require the generation of empirical null distributions derived from the periodicity and regulator metrics of many synthetic expression profiles generated by repeated sampling of the experimental data. As the number of genes and/or the number of time points increases, the background distributions of potential random synthetic abundance profiles grows rapidly. As a result, in general, many more synthetic profiles must be generated and characterized to improve estimates of these *p*-values. If too few random curves are analyzed, there may be ambiguity in the final rankings due to repeated *p*-values caused by the resulting coarse discretization of possible estimates. Additionally, the choice of a background distribution has a large impact on statistical significance [27] and gives poor results when assumptions of the background distribution do not match the reality of the data (see the discussion of the malaria data set in [28]).

Is it necessary to compute a significance value in order to skillfully rank core TFs? We address this question by ranking genes according to DL’s “naive” measurements for periodicity and regulation, individually (DL Per Score and DL Reg Score in Table 1, respectively) and in combination (PerReg). These naive measurements are calculated quickly with no permutations or random sampling required, and thus greatly reduce the computational time required to rank genes. When used individually, the naive DL measurements perform equally well or better than the empirical *p*-values at identifying core, as measured by AP (Fig. 2B). Indeed, there is a striking difference across all datasets in the ranking of core genes using DL’s naive periodicity score rather than its associated empirical *p*-value, which is particularly expensive to compute for large gene sets.

When combined, the naive measures also skillfully rank genes well above baseline across all datasets. In fact, there is a notable increase in AP over the other combined metrics, which are derived from *p*-values, for the *A. thaliana* data in both conditions (Fig. 2A). We expect that this, along with the generally lower performance of these metrics on *A. thaliana* data compared to the other datasets, may be due to the fact that the *A. thaliana* transcript abundance profiles reflect gene expression in multiple tissue types, making it difficult to collect accurate empirical *p*-values.

Much like DL × JTK, PerReg shows skillful recall at identifying core genes among the top 25 TFs (Table S3), identifying at least 4 and at most 10 core TFs among the top 25, across all datasets considered in this study.

### Several high ranking non-core genes display regulatory relationships with core genes

The lists of core TFs used in this study are conservative since 1) a lack of strong evidence supporting a gene as a core regulator is not proof that it is not core and 2) many functional regulators are also known to be transcriptional co-regulators and post-transcriptional modifiers; we labelled the latter as non-core to ensure fair assessment of the performance of the ranking methods. Thus, our binary labels may contain false negatives (core labeled as non-core) due to a lack of strong experimental evidence, and certainly contain false negatives due to our restriction to TFs. We ask, what are the identities of the most highly ranked non-core TFs, and does there exist any evidence that they target the activity of and/or are targeted by our core TFs?

Utilizing the curated list of regulatory relationships in YEASTRACT [22] and PlantTFDB [21], as well as a literature search for *M. musculus* TF interactions, we indeed observe evidence that several yeast, plant, and mouse genes among the top 25 TFs ranked by the measure DL × JTK target core and/or are targeted by core (Table 4). For example, we find that among the top 25 *S. cerevisiae* TFs ranked by DL × JTK in either MA or RNASeq datasets, that 40% (9/23) of the genes have existing evidence of both regulating and being regulated by core. This observation suggests that genes that appear highly ranked by our combined measures, but were not labeled as core due to a lack of existing evidence, may in fact be core nodes.

**Table 4.** Interaction relationships*\** between core TFs and non-core that appear in the top 25 TFs as ranked by DL×JTK^*†*^.

| *S. cerevisiae* |  |  | *M. musculus* |  |  | *A. thaliana* |  |  |
| --- | --- | --- | --- | --- | --- | --- | --- | --- |
| Gene | Targeted by | Targets | Gene | Targeted by | Targets | Gene | Targeted by | Targets |
| *ASG1* | FHL1 | *NDD1* | *EGR1* | ARNTL [29] | *ARNTL* [30] | *EPR1* | RVE4 | *PRR5* |
| *EDS1* | FHL1 | *TOS4* | *KLF10* | ARNTL [31] | *ARNTL* [32] | *PIF4* | CCA1 | *LHY* |
| *GAT1* | ACE2 | *ACE2* | *NFIC* | HLF [33] |  | *PIL6* | CCA1 | *LHY* |
| *MTH1* | FHL1 | *STB1* | *ATF5* | CLOCK [34] |  | *ARF11* | CHE |  |
| *RME1* | ACE2 | *ASH1* | *ESR1* | CLOCK [35] |  | *CO* | CCA1 |  |
| *RPI1* | FHL1 | *NDD1* | *SREBF1* |  | *BHLHE40/41* [36] | *COL1* | CHE |  |
| *SIP4* | FHL1 | *STB1* |  |  |  | *COL9* | CHE |  |
| *TEC1* | SWI4 | *ASH1* |  |  |  | *MYBL2* | CHE |  |
| *WTM2* | ACE2 | *STB1* |  |  |  | *CDF2* |  | *LHY* |
| *ASF1* | SWI4 |  |  |  |  | *RVE1* |  | *PRR5* |
| *HST3* | FKH1 |  |  |  |  | *RVE2* |  | *CCA1* |
| *HST4* | MBP1 |  |  |  |  |  |  |  |
| *POG1* | MCM1 |  |  |  |  |  |  |  |
| *RLF2* | MBP1 |  |  |  |  |  |  |  |
| *RTT107* | MCM1 |  |  |  |  |  |  |  |
| *SNF5* | ACE2 |  |  |  |  |  |  |  |
| *TBF1* | FHL1 |  |  |  |  |  |  |  |

\* *S. cerevisiae* and *A. thaliana* interactions determined respectively by database searches of [22] and [21] and represent a range of direct and indirect evidence types, including the presence of binding motifs in regulatory regions and response to TF over-expression. *M. musculus* interactions determined by evidence gathered in the associated citation.
† *M. musculus* non-core TFs drawn from MA dataset only, while non-core *S. cerevisiae* and *A. thaliana* TFs were drawn from the unions of each pair of analyzed datasets.

Within the top 25 of all genes, as ranked by DL×JTK, we observe a number of regulatory elements that are known to be essential to produce the given periodic program of gene expression, but which are not strictly TFs, and therefore do not qualify in our definition as a core gene. Examples include the mouse transcriptional co-regulators Period 3 (PER3) [37] and Cryptochrome 1 (CRY1) [38] and the plant post-transcriptional gene Gigantea (GI) [39] (Table 4), which are known or proposed to be transcriptional co-regulators and post-transcriptional elements. This supports our conclusion that core elements, even beyond the TFs, can be identified by quantifiable features in their transcript abundance dynamics. Improvement in the annotation of non-TF regulatory elements is needed before we can reliably quantify the extent to which these dynamic characteristics are exhibited by all nodes of these networks at the level of transcript abundance.

### External periodic signals do not significantly alter the skill of transcript abundance dynamics at identifying core genes

Implicit in the definitions of the core transcriptional regulatory networks considered in this study is that they are free-running and can support rhythmic oscillations in the absence of external periodic stimuli due to their mutual regulatory interactions with other core elements. Is necessary to collect time series transcriptomics in the absence of external circadian stimuli to skillfully identify core regulatory elements?

To address this question, we compared the skill of dynamic expression features to identify the core TFs for *A. thaliana* in 1) periodically fluctuating light and temperature (diurnal) conditions (LDHC) and 2) constant light, (circadian) conditions (LL LDHC). For the details on the precise experimental setup see [14].

One might expect that the transcript dynamics of diurnal non-core genes—those that are strictly driven by periodic light-dark and/or temperature cycles—would reduce the capacity of dynamic gene expression features to distinguish core regulatory elements. We find that the signal of core genes is not degraded in the presence of external periodic stimuli in these experiments, since all combined quantitative measures show nearly identical skill at identifying core genes across both conditions (Fig. 2A). Even more striking is the consistency in the individual ranks of core genes across diurnal and circadian conditions, as shown for DL×JTK in Table S1.

## Conclusion

Elucidating the underlying GRNs driving dynamic biological processes, such as cell-division and sleep-wake cycles, is crucial if we are to leverage existing control mechanisms for synthetic biology applications, understand the evolution of biological networks, and inform experiments to discover new drug targets. However, experimentally identifying the core regulatory elements of these gene networks can be costly, time consuming and daunting, even for the simplest organisms, due to the large hypothesis space. We have shown that many core transcriptional regulators, appearing in organisms separated by millions of years of evolution, share common features in their transcript abundance dynamics. We demonstrated the use of several metrics that quantify and combine these dynamic features. The outcome is a substantial reduction in hypothesis space, a prioritization of gene targets for experimental validation, and a facilitation of network modeling via the identification of control variables.

High degrees of periodicity and strong regulation signals appear to be characteristic features of many core TFs involved in generating periodic biological processes. However, not all known core regulatory TFs strongly exhibit the dynamic features quantified here at the level of their transcript abundance. For instance, the abundance profile of the core *S. cerevisiae* TF *NDD1* is highly periodic with a precise match to cell-cycle period and exhibits large dynamic range, but *MCM1* does not show convincing oscillations at the cell-cycle period (Fig. 3A). *MCM1* is the only core TF to not rank in the top 70 TFs in at least one of the two *S. cerevisiae* datasets using DL × JTK (Table S1). However, MCM1 acts in complex with other rhythmically-expressed genes like NDD1 [40, 41], so it can still be part of a highly periodic TF complex without itself exhibiting highly periodic signatures in transcript abundance. It is enticing to imagine there may be other features captured in the gene or protein expression profiles, as well as features not related to gene expression, such as sequence-based and protein interaction features that could be used to more accurately capture all core genes, including those identified in TF complexes.

It is known in the circadian field that several core clock genes have tissue-specific periodic properties in mice [13]. Thus, we expect not all core genes will rise to the top of our rankings in every tissue. For example, within the three retinoid-related orphan receptors (RORs) TFs, *RORA, RORB*, and *RORC*, only *RORC* is known to display periodic gene expression in mouse liver [42]. Indeed, only *RORC* was ranked in the top 25 TFs ranked by DL × JTK (Table 3) in the mouse liver microarray dataset. Another example is the mouse core clock gene *ARNTL2*, which is not ranked highly in the mouse liver datasets. Most studies suggest *ARNTL2* has brain-specific circadian expression with lower levels of expression in the liver in mammals [43–45]. There is also growing evidence for genes to exhibit tissue-specific dynamics in plants [46].

Our ability to identify plant core genes appears generally lower than the other organisms we considered. This may be due to the fact that samples were taken from the whole leaf and thus contained a mixture of multiple tissue types such as mesophyll, epidermis, and vasculature [14]. The abundance and periodicity of any particular transcript might therefore appear muted as genes are likely expressed differentially across tissues. Consistent with this hypothesis, several studies have shown that tissue-specific clocks in plants can be asymmetrically coupled [47], have different period lengths [48], or have different levels of gene expression for core components [49, 50]. Naturally it is more difficult to identify a core component whose observed dynamics is either a convolution of multiple dissimilar abundance profiles derived from multiple tissues or has specificity to an under-represented tissue in a mixture of tissue types. Interestingly, the dominant tissue type in whole leaf samples is mesophyll, and morning-expressed clock genes (*CCA1*, PRRs, and *LHY*) are highly expressed in the mesophyll [47, 51]. These morning-expressed genes are mostly the only plant core genes ranked highly in this study (Table 3).

Broadly speaking, our findings herein suggest that even naive measures of periodicity and regulatory strength can be used to skillfully rank genes. We conclude that classifiers are likely dependent on the dynamic characteristics of the the transcript abundance profiles, and perhaps less so than on the particular quantification of these characteristics. Thus, broad recommendations for thresholds that reliably identify core nodes are currently not possible. That said, with the availability of proper experimental controls across organism, platform, sampling density, etc., it might be possible to compare the various metrics to make a more prescriptive recommendation of which particular method to use for a given dataset.

The use of naive metrics rather than empirical *p*-values does not suffer from ambiguous rankings caused by insufficient sampling of the null distribution, as may be the case with DL’s method of measuring significance. It is possible to reduce the ambiguity of a ranking by increasing the sampling of the null distribution at the cost of increased compute time. The disambiguation of empirical regulator *p*-values computed by the DL metric through increased sampling is visualized in Fig. S7. Similarly, combining several *p*-values derived from different dynamic characteristics into combined metrics can eliminate ambiguous rankings that may be present in one of these features.

We have demonstrated the importance of reliable genome annotation of TF genes, but many organisms of interest currently lack comprehensive gene annotations. Thus it is desirable to have methods that can leverage high-throughput technologies to provide evidence of gene function. Additional evidence such as identifying DNA-binding domains and/or orthology to known TFs in other organisms are two such methods that could be used to provide putative TF lists for poorly-annotated genomes.

Here we demonstrate that dynamic features of transcriptomes appear to be conserved across kingdoms and networks that appear to serve disparate functions such as cell-cycle or circadian clocks. It is possible that the conservation of these features results from a fundamental property of GRNs, where a transcriptional signal is developed within a core set of nodes and that the signal degrades as it is propagated through effector nodes that control downstream gene expression. Alternatively, the conservation of features could reflect an evolutionary conservation of network topologies that produce rhythmic behaviors during circadian and cell cycles.

## Supporting information

Dataset S1: Gene Expression Data

Dataset S2: Core Genes

Dataset S3: Transcription Factors

Dataset S4: Gene Rankings

## Supporting Information

**Fig. S1**. Precision-recall curves of classifiers identifying core from non-core genes in the *S. cerevisiae* microarray dataset.

**Fig. S2**. Precision-recall curves of classifiers identifying core from non-core genes in the *S. cerevisiae* RNASeq dataset.

**Fig. S3**. Precision-recall curves of classifiers identifying core from non-core genes in the *M. Musculus* microarray dataset.

**Fig. S4**. Precision-recall curves of classifiers identifying core from non-core genes in the *M. Musculus* RNASeq dataset.

**Fig. S5**. Precision-recall curves of classifiers identifying core from non-core genes in the *A. thaliana* microarray, diurnal condition dataset.

**Fig. S6**. Precision-recall curves of classifiers identifying core from non-core genes in the *A. thaliana* microarray, circadian condition dataset.

**Fig. S7**. Plot of the number of unique DL Reg *p*-values as a function of the number of permutations used in the calculation.

**Table S1** Ranks of all core genes among transcription factors using DL*×*JTK score.

**Table S2** Top 25 transcription factors, ranked by DL score.

**Table S3** Top 25 transcription factors, ranked by PerReg score.

**Dataset S1 Gene Expression Data**. An EXCEL file containing gene expression profiles for each dataset used in this study.

**Dataset S2 Core Genes**. An EXCEL file containing lists of core genes for all organisms.

**Dataset S3 Transcription Factors**. An EXCEL file containing lists of transcription factors for all organisms.

**Dataset S4 Gene Rankings**. An EXCEL file containing the rankings of all genes by each metric for all datasets.

## Code and Data Availability

All data and code used to process and analyze the data, and generate figures are provided in a public repository at [52].

## References

1. Harmer SL. The Circadian System in Higher Plants. Annual Review of Plant Biology. 2009;60(1):357–377. doi:10.1146/annurev.arplant.043008.092054.

2. Brunner M, Schafmeier T. Transcriptional and post-transcriptional regulation of the circadian clock of cyanobacteria and Neurospora. Genes and Development. 2006;20:1061–1074. doi:10.1101/gad.1410406.

3. Panda S, Hogenesch J, Kay S. Circadian rhythms from flies to human. Nature. 2002;417:329–335. doi:10.1038/417329a.

4. Bristow SL, Leman AR, Kovacs LAS, Deckard A, Harer J, Haase SB. Checkpoints couple transcription network oscillator dynamics to cell-cycle progression. Genome Biology. 2014;15(9):446. doi:https://doi.org/10.1186/s13059-014-0446-7.

5. Simmons Kovacs L, Mayhew M, Orlando D, Jin Y, Li Q, Huang C, et al. Cyclin-Dependent Kinases Are Regulators and Effectors of Oscillations Driven by a Transcription Factor Network. Molecular Cell. 2012;45(5):669–679. doi:https://doi.org/10.1016/j.molcel.2011.12.033.

6. Orlando DA, Lin CY, Bernard A, Wang JY, Socolar JE, Iversen ES, et al. Global control of cell-cycle transcription by coupled CDK and network oscillators. Nature. 2008;453(7197):944.

7. Haase SB, Reed SI. Evidence that a free-running oscillator drives G1 events in the budding yeast cell cycle. Nature. 1999;401:394–397. doi:10.1038/43927.

8. Cho CY, Kelliher CM, Haase SB. The cell-cycle transcriptional network generates and transmits a pulse of transcription once each cell cycle. Cell Cycle. 2019;18(4):363–378.

9. Hughes ME, Hogenesch JB, Kornacker K. JTK CYCLE: An Efficient Nonparametric Algorithm for Detecting Rhythmic Components in Genome-Scale Data Sets. Journal of Biological Rhythms. 2010;25(5):372–380. doi:10.1177/0748730410379711.

10. de Lichtenberg U, Jensen LJ, Fausbøll A, Jensen TS, Bork P, Brunak S. Comparison of computational methods for the identification of cell cycle-regulated genes. Bioinformatics. 2005;21(7):1164–1171. doi:10.1093/bioinformatics/bti093.

11. Saito T, Rehmsmeier M. The Precision-Recall Plot Is More Informative than the ROC Plot When Evaluating Binary Classifiers on Imbalanced Datasets. PLOS ONE. 2015;10(3):1–21. doi:10.1371/journal.pone.0118432.

12. Kelliher CM, Leman AR, Sierra CS, Haase SB. Investigating Conservation of the Cell-Cycle-Regulated Transcriptional Program in the Fungal Pathogen, Cryptococcus neoformans. PLOS Genetics. 2016;12(12):1–23. doi:10.1371/journal.pgen.1006453.

13. Zhang R, Lahens NF, Ballance HI, Hughes ME, Hogenesch JB. A circadian gene expression atlas in mammals: Implications for biology and medicine. Proceedings of the National Academy of Sciences. 2014;111(45):16219–16224. doi:10.1073/pnas.1408886111.

14. Mockler TC, Michael TP, Priest HD, Chen R, Sullivan CM, Givan SA, et al. The Diurnal Project: Diurnal and Circadian Expression Profiling, Model-based Pattern Matching, and Promoter Analysis. Cold Spring Harbor Symposia on Quantitative Biology. 2007;72:353–363. doi:10.1101/sqb.2007.72.006.

15. Orlando D, Lin CY, Bernard A, Iversen ES, Hartemink AJ, Haase SB. A Probabilistic Model for Cell Cycle Distributions in Synchrony Experiments. Cell Cycle. 2007;6(4):478–488. doi:10.4161/cc.6.4.3859.

16. McGoff KA, Guo X, Deckard A, Kelliher CM, Leman AR, Francey LJ, et al. The Local Edge Machine: inference of dynamic models of gene regulation. Genome Biology. 2016;17(1):214. doi:10.1186/s13059-016-1076-z.

17. Lowrey PL, Takahashi JS. MAMMALIAN CIRCADIAN BIOLOGY: Elucidating Genome-Wide Levels of Temporal Organization. Annual Review of Genomics and Human Genetics. 2004;5(1):407–441. doi:10.1146/annurev.genom.5.061903.175925.

18. Takahashi J. Transcriptional architecture of the mammalian circadian clock. Nature Review Genetics. 2017;18:167 – 197. doi: https://doi.org/10.1038/nrg.2016.150.

19. Haase SB, Wittenberg C. Topology and Control of the Cell-Cycle-Regulated Transcriptional Circuitry. Genetics. 2014;196(1):65–90. doi:10.1534/genetics.113.152595.

20. Hu H, Miao YR, Jia LH, Yu QY, Zhang Q, Guo AY. AnimalTFDB 3.0: a comprehensive resource for annotation and prediction of animal transcription factors. Nucleic Acids Research. 2018;47(D1):D33–D38. doi:10.1093/nar/gky822.

21. Jin J, Tain F, Yang DC, Meng YQ, Kong L, Luo J, et al. PlantTFDB 4.0: toward a central hub for transcription factors and regulatory interactions in plants. Nucleic Acids Research. 2016;45(D1):D1040–D1045. doi:10.1093/nar/gkw982.

22. Teixeira MC, Monteiro PT, Palma M, Costa C, Godinho CP, Pais P, et al. YEASTRACT: an upgraded database for the analysis of transcription regulatory networks in Saccharomyces cerevisiae. Nucleic Acids Research. 2017;46(D1):D348–D353. doi:10.1093/nar/gkx842.

23. Nakamichi N, Kiba T, Henriques R, Mizuno T, Chua NH, Sakakibara H. PSEUDO-RESPONSE REGULATORS 9, 7, and 5 Are Transcriptional Repressors in the Arabidopsis Circadian Clock. The Plant Cell. 2010;22(3):594–605. doi:10.1105/tpc.109.072892.

24. Kim H, Kim HJ, Vu QT, Jung S, McClung CR, Hong S, et al. Circadian control of ORE1 by PRR9 positively regulates leaf senescence in Arabidopsis. Proceedings of the National Academy of Sciences. 2018;115(33):8448–8453. doi:10.1073/pnas.1722407115.

25. Gendron JM, Pruneda-Paz JL, Doherty CJ, Gross AM, Kang SE, Kay SA. Arabidopsis circadian clock protein, TOC1, is a DNA-binding transcription factor. Proceedings of the National Academy of Sciences. 2012;109(8):3167–3172. doi:10.1073/pnas.1200355109.

26. Deckard A, Anafi RC, Hogenesch JB, Haase SB, Harer J. Design and analysis of large-scale biological rhythm studies: a comparison of algorithms for detecting periodic signals in biological data. Bioinformatics. 2013;29(24):3174–3180. doi:10.1093/bioinformatics/btt541.

27. Futschik ME, Herzel H. Are we overestimating the number of cell-cycling genes? The impact of background models on time-series analysis. Bioinformatics. 2008;24(8):1063–1069. doi:10.1093/bioinformatics/btn072.

28. Kallio A, Vuokko N, Ojala M, Haiminen N, Mannila H. Randomization techniques for assessing the significance of gene periodicity results. BMC Bioinformatics. 2011;12(1):330. doi:10.1186/1471-2105-12-330.

29. Lee JH, Sancar A. Circadian clock disruption improves the efficacy of chemotherapy through p73-mediated apoptosis. Proceedings of the National Academy of Sciences. 2011;108(26):10668–10672. doi:10.1073/pnas.1106284108.

30. Riedel CS, Georg B, Jørgensen HL, Hannibal J, Fahrenkrug J. Mice Lacking EGR1 Have Impaired Clock Gene (BMAL1) Oscillation, Locomotor Activity, and Body Temperature. Journal of molecular neuroscience : MN. 2018;64(1):9—19. doi:10.1007/s12031-017-0996-8.

31. Guillaumond F, Gréchez-Cassiau A, Subramaniam M, Brangolo S, Peteri-Brünback B, Staels B, et al. Krüppel-Like Factor KLF10 Is a Link between the Circadian Clock and Metabolism in Liver. Molecular and Cellular Biology. 2010;30(12):3059–3070. doi:10.1128/MCB.01141-09.

32. Hirota T, Kon N, Itagaki T, Hoshina N, Okano T, Fukada Y. Transcriptional repressor TIEG1 regulates Bmal1 gene through GC box and controls circadian clockwork. Genes to Cells. 2010;15(2):111–121. doi:10.1111/j.1365-2443.2009.01371.x.

33. Wahlestedt M, Ladopoulos V, Hidalgo I, Castillo MS, Hannah R, Säwén P, et al. Critical Modulation of Hematopoietic Lineage Fate by Hepatic Leukemia Factor. Cell Reports. 2017;21(8):2251 – 2263. doi: https://doi.org/10.1016/j.celrep.2017.10.112.

34. Lemos DR, Goodspeed L, Tonelli L, Antoch MP, Ojeda SR, Urbanski HF. Evidence for Circadian Regulation of Activating Transcription Factor 5 But Not Tyrosine Hydroxylase by the Chromaffin Cell Clock. Endocrinology. 2007;148(12):5811–5821.

35. Yoshitane H, Ozaki H, Terajima H, Du NH, Suzuki Y, Fujimori T, et al. CLOCK-Controlled Polyphonic Regulation of Circadian Rhythms through Canonical and Noncanonical E-Boxes. Molecular and Cellular Biology. 2014;34(10):1776–1787. doi:10.1128/MCB.01465-13.

36. Lecomte V, Meugnier E, Euthine V, Durand C, Freyssenet D, Nemoz G, et al. A New Role for Sterol Regulatory Element Binding Protein 1 Transcription Factors in the Regulation of Muscle Mass and Muscle Cell Differentiation. Molecular and Cellular Biology. 2010;30(5):1182–1198. doi:10.1128/MCB.00690-09.

37. Zhang L, Hirano A, Hsu PK, Jones CR, Sakai N, Okuro M, et al. A PERIOD3 variant causes a circadian phenotype and is associated with a seasonal mood trait. Proceedings of the National Academy of Sciences of the United States of America. 2016;113(11):E1536–E1544. doi:10.1073/pnas.1600039113.

38. van der Horst GTJ, Muijtjens M, Kobayashi K, Takano R, ichiro Kanno S, Takao M, et al. Mammalian Cry1 and Cry2 are essential for maintenance of circadian rhythms. Nature. 1999;398:627–630. doi:10.1038/19323.

39. Mishra P, Panigrahi KC. GIGANTEA – an emerging story. Frontiers in Plant Science. 2015;6:8. doi:10.3389/fpls.2015.00008.

40. Kelliher CM, Foster MW, Motta FC, Deckard A, Soderblom EJ, Moseley MA, et al. Layers of regulation of cell-cycle gene expression in the budding yeast Saccharomyces cerevisiae. Molecular Biology of the Cell. 2018;29(22):2644–2655. doi:10.1091/mbc.E18-04-0255.

41. Koranda M, Schleiffer A, Endler L, Ammerer G. Forkhead-like transcription factors recruit Ndd1 to the chromatin of G2/M-specific promoters. Nature. 2000;406:94–98. doi:doi.org/10.1038/35017589.

42. Ueda HR, Chen W, Adachi A, Wakamatsu H, Hayashi S, Takasugi T, et al. A transcription factor response element for gene expression during circadian night. Nature. 2002;418:534–539. doi:10.1038/nature00906.

43. Ikeda M, Yu W, Hirai M, Ebisawa T, Honma S, Yoshimura K, et al. cDNA Cloning of a Novel bHLH-PAS Transcription Factor Superfamily Gene, BMAL2: Its mRNA Expression, Subcellular Distribution, and Chromosomal Localization. Biochemical and Biophysical Research Communications. 2000;275(2):493 – 502. doi: https://doi.org/10.1006/bbrc.2000.3248.

44. Maemura K, de la Monte SM, Chin MT, Layne MD, Hsieh CM, Yet SF, et al. CLIF, a Novel Cycle-like Factor, Regulates the Circadian Oscillation of Plasminogen Activator Inhibitor-1 Gene Expression. Journal of Biological Chemistry. 2000;275(47):36847–36851. doi:10.1074/jbc.C000629200.

45. Hogenesch JB, Gu YZ, Moran SM, Shimomura K, Radcliffe LA, Takahashi JS, et al. The Basic Helix-Loop-Helix-PAS Protein MOP9 Is a Brain-Specific Heterodimeric Partner of Circadian and Hypoxia Factors. Journal of Neuroscience. 2000;20(13):RC83–RC83. doi:10.1523/JNEUROSCI.20-13-j0002.2000.

46. Inoue K, Araki T, Endo M. Oscillator networks with tissue-specific circadian clocks in plants. Seminars in Cell and Development Biology. 2018;83:78–85. doi:10.1016/j.semcdb.2017.09.002.

47. Endo M, Shimizu H, Nohales MA, Araki T, Kay SA. Tissue-specific clocks in Arabidopsis show asymmetric coupling. Nature. 2014;515:419–422. doi:10.1038/nature13919.

48. Yakir E, Hassidim M, Melamed-Book N, Hillman D, Kron I, Green RM. Cell autonomous and cell-type specific circadian rhythms in Arabidopsis. The Plant Journal. 2011;68(3):520–531.

49. Para A, Farré EM, Imaizumi T, Pruneda-Paz JL, Harmon FG, Kay SA. PRR3 Is a Vascular Regulator of TOC1 Stability in the Arabidopsis Circadian Clock. The Plant Cell. 2007;19(11):3462–3473. doi:10.1105/tpc.107.054775.

50. Edwards J, Martin AP, Andriunas F, Offler CE, Patrick JW, McCurdy DW. GIGANTEA is a component of a regulatory pathway determining wall ingrowth deposition in phloem parenchyma transfer cells of Arabidopsis thaliana. Plant Journal. 2010;63(4):651–661.

51. Endo M, Shimizu H, Araki T. Rapid and simple isolation of vascular, epidermal and mesophyll cells from plant leaf tissue. Nature Protocols. 2016;11:1388–1395. doi:10.1038/nprot.2016.083.

52. Moseley RC. Dynamic Feature Analysis; 2020. https://gitlab.com/bertfordley/dynamic_features_analysis.

