## Supplemental Methods, Figures, & Tables for "Conservation of dynamic characteristics of transcriptional regulatory elements in periodic biological processes"

### 1 Methods

#### Dynamic Curve Features

##### De Lichtenberg Measures

In [1] the authors introduced a periodicity detection algorithm designed to identify genes in yeast that oscillated with the cell division cycle. The de Lichtenberg algorithm (DL) measures how periodic a signal is at a specified period by quantifying and combining statistical measures of gene expression periodicity and strength of regulation (Eq. 1). For each gene expression profile two empirical $p$-values, $p_{reg}$ and $p_{per}$, are independently computed. Respectively, these $p$-values estimate the probabilities that the observed fold-change variability (Eq. 2) and rhythmicity at a specified period (Eq. 3) of the expression profile occurred, in some sense, at random. These statistics are then combined in a manner which accentuates expression profiles that are simultaneously highly-periodic and highly-variable. Explicitly, let $G \in \mathcal{G}$ be the gene expression profile, measured at timepoints $T = (t_1, \ldots, t_n)$, corresponding to gene $G$ in the set of all measured gene expression profiles in a collection, $\mathcal{G}$. Then

$$\mathrm{DL}(G) := p_{\mathrm{reg}}(G) p_{\mathrm{per}}(G) \left[1 + \left(\frac{p_{\mathrm{reg}}(G)}{0.001}\right)^2\right] \left[1 + \left(\frac{p_{\mathrm{per}}(G)}{0.001}\right)^2\right]. \tag{1}$$

The $p$-value $p_{\mathrm{reg}}(G)$ is meant to be the probability that a "random" curve appears more highly regulated than $G$. This variability metric, denoted $\mathrm{Reg}(G)$, and defined by

$$\mathrm{Reg}(G) = \mathrm{std}\left(\log_{10}\left(G/\mathrm{mean}(G)\right)\right) \tag{2}$$

captures the deviation of the time series about its mean, with a small value indicating little variation over time about the mean expression level. Thus, $\mathrm{Reg}(G)$ may be interpreted as the magnitude of regulation of the gene $G$. The rationale of this study is rooted in the expectation that $\mathrm{Reg}(G)$ will actually be largest among those genes which are primarily responsible for generating an observed program of dynamic transcript abundance.

In practice, to estimate $p_{\text{reg}}(G)$, an empirical null distribution of curve variability metrics is generated by first creating a large number of random expression curves through selection at each experimental time point the expression value of a transcript chosen from $\mathcal{G}$ uniformly at random. The variability metric (Reg) of each random curve is then computed and $p_{\text{reg}}(G)$ is taken as the fraction of random curves whose regulation score is larger than $\text{Reg}(G)$.

Likewise, for each gene $G$, many simulated curves are generated by randomly permuting the expression values of $G$ and comparing the periodicity metric (called a Fourier score in [1]) of the randomized curve to that of $G$. The $p$-value $p_{\text{per}}(G)$ is the fraction of simulated curves whose periodicity score (Per) is larger than that of $G$. This periodicity metric is taken to be

$$\text{Per}(G) := \sqrt{(G \cdot \cos(\omega T))^2 + (G \cdot \sin(\omega T))^2}, \quad (3)$$

where $\omega$ is a specified period, and therefore reflects the magnitude of the Fourier coefficient at that period. A new implementation of the DL algorithm was written for use in this analysis and is available as a Python 3 module under the MIT License [2].

##### JTK-CYCLE Measures

JTK-CYCLE (JTK) was developed as a periodicity-detection algorithm to identify circadian rhythmically expressed genes in mice [3]. Since then it has been successfully applied to time series transcriptomics data collected from many other species exhibiting rhythmic phenotypes. JTK correlates a gene's expression profile to that of a reference curve with known periodicity properties, and computes a significance of that correlation. Usually, and in this analysis, sinusoidal template curves are generated with user-specified periods and at various phase shifts determined by the sampling times of the expression profiles. A pattern of "ups" and "downs" is computed—by comparing the expression level at each time with all subsequent times—for both the expression curve $G$ and the equivalently sampled sinusoidal template curves. Then the total number of agreements (concordances) and disagreements (discordances) in the up-down pattern of $G$ and that of the known periodic curves are computed, giving the Kendall rank correlation coefficient between the curves. By precomputing the exact null distribution of Kendall's tau correlation [4] using the Harding algorithm [5], an exact Bonferroni-adjusted $p$-value is rapidly computed for each gene, $G$, and is denoted here by $p_{jtk}(G)$. An implementation of JTK in Python by Alan Hutchinson [6] was modified for use in this analysis and is available as a Python 3 module under the MIT License [7].

#### Precision-Recall Curves & Average Precision

Given a ranking of $N$ genes by some metric that is meant to discriminate between core and non-core, for some choice of threshold, we examine the true positives (TP) compared to false positives (FP) and/or false negatives (FN). The *precision* of the classifier's ranking at a given threshold is the fraction of true core genes among all genes ranked above that threshold, i.e. TP / (TP + FP). The *recall* is the fraction of true core genes appearing above the chosen threshold: TP / (TP + FN). Any ranking can achieve a perfect recall of 1 if the threshold is chosen so permissively as to label all genes as core. However, given the goal to reduce hypothesis space and limit the amount of experimental validation needed to identify core regulatory elements, a permissive choice of threshold is of little practical utility. Thus, in this context, a meaningful measure of classifier skill is the precision at a given recall. For example, the precision at a recall of

10 percent characterizes how many knock-out experiments one would expect to perform, in accordance with a given algorithm's ranking of genes, before 10 percent of the core regulatory elements are identified. It is this interpretation that may be of particular value to a researcher interested in using a ranking algorithm to prioritize experiments.

In some rare cases, if a scoring algorithm is particularly discriminating between two classes, the scores may be bimodally distributed and well-separated, allowing a data-driven justification of a threshold. Usually, this is not the case, and a threshold must be chosen arbitrarily. Thus, better measures of classifier performance incorporate algorithm performance across all thresholds. These measures assess the ranking itself, quantifying the skill of the classifier to rank the members of the true class (core) above the members of the other class (non-core). One such measure is the so-called *average precision* (AP) of the *precision-recall* (PR) curve defined to be $\sum_{n=1}^{N} P_n \Delta R_n$, where $\Delta R_n$ is the change in the recall level caused by moving the decision threshold from the $(n-1)$st gene to the $n$th gene in the list of genes ranked by some metric, and $P_n$ is the precision of the metric when the threshold is set to the metric of the $n$th gene in the ranked list.

### 2 Tables

**Table S1. Ranks among all transcription factors of core genes using the DL×JTK score.**

| *S. cerevisiae* | | | *M. Musculus* | | | *A. thaliana* | | |
|---|---|---|---|---|---|---|---|---|
| Gene | MA | RNA | Gene | MA | RNA | Gene | LDHC | LL_LDHC |
| *TOS4* | 16 | 1 | *DBP* | 2 | 1 | *CCA1* | 11 | 3 |
| *SWI5* | 1 | 4 | *NPAS2* | 3 | 2 | *LHY* | 6 | 11 |
| *YOX1* | 2 | 5 | *ARNTL* | 1 | 4 | *LUX* | 30 | 16 |
| *ASH1* | 14 | 8 | *NR1D1* | 4 | 20 | *PRR7* | 12 | 17 |
| *FKH1* | 10 | 9 | *NR1D2* | 5 | 45 | *RVE8* | 15 | 21 |
| *ACE2* | 5 | 11 | *RORC* | 10 | 67 | *PRR9* | 49 | 30 |
| *SWI4* | 12 | 13 | *NFIL3* | 8 | 71 | *TOC1* | 44 | 34 |
| *PLM2* | 11 | 15 | *TEF* | 11 | 87 | *PRR5* | 48 | 45 |
| *NDD1* | 13 | 17 | *BHLHE41* | 6 | 123 | *CHE* | 97 | 138 |
| *HCM1* | 8 | 18 | *HLF* | 69 | 164 | *RVE4* | 63 | 222 |
| *STB1* | 7 | 21 | *CLOCK* | 7 | 419 | *RVE6* | 453 | 908 |
| *YHP1* | 15 | 22 | *BHLHE40* | 195 | 501 | | | |
| *FKH2* | 49 | 28 | *ARNTL2* | 982 | 671 | | | |
| *FHL1* | 20 | 40 | *RORA* | 283 | 823 | | | |
| *SWI6* | 176 | 50 | *RORB* | 1210 | NA | | | |
| *MBP1* | 151 | 66 | | | | | | |
| *MCM1* | 266 | 146 | | | | | | |
| **No. of TFs**† | 304 | 307 | | 1373 | 1118 | | 1415 | 1415 |

LL_LDHC: Constant light and temperature; LDHC: 24 hour cycling light and temperature; MA: Microarray; RNA: RNAseq

† Counts are based on post-processed datasets (see Materials and Methods)

**Table S2. The top 25 highest DL-ranked genes among all transcription factors.**

| Rank | *S. cerevisiae* MA | *S. cerevisiae* RNA | *M. Musculus* MA | *M. Musculus* RNA | *A. thaliana* LDHC | *A. thaliana* LL_LDHC |
|---|---|---|---|---|---|---|
| 1 | ASH1* | RME1 | ARNTL* | NPAS2* | LHY* | RVE1 |
| 2 | RME1 | YOX1* | DBP* | DBP* | BBX19 | PIF4 |
| 3 | SWI5* | ASH1* | NPAS2* | CDX4 | COL2 | BBX18 |
| 4 | HST4 | TOS4* | NR1D1* | GM14444 | RVE8* | COL5 |
| 5 | NUT1 | KAR4 | BHLHE41* | ARNTL* | TOC1* | LHY* |
| 6 | HST3 | RTT107 | NR1D2* | NR1D1* | COL1 | STH |
| 7 | CIN5 | SWI5* | NFIL3* | CREB5 | COL9 | PRR7* |
| 8 | ACE2* | SWI4* | EGR1 | PPARD | STH | RVE8* |
| 9 | MET4 | TEC1 | TEF* | NPAS3 | TGA3 | COL1 |
| 10 | SWI4* | HST4 | PPARD | POU4F1 | RVE1 | COL2 |
| 11 | YOX1* | ASF1 | RORC* | FOXO6 | HYH | AT2G28200 |
| 12 | ISW2 | HST3 | MAFB | DMRTA2 | BBX18 | PIL6 |
| 13 | RGT2 | RLF2 | KLF10 | EGR1 | AT1G26790 | MYBL2 |
| 14 | MET18 | WTM1 | RFXANK | GM20422 | PRR7* | CCA1* |
| 15 | RGT1 | ACE2* | CLOCK* | EGR3 | HB-12 | BBX8 |
| 16 | SIP4 | ZNF1 | TSC22D3 | TBX1 | PRE1 | HSFA8 |
| 17 | TOS4* | STB1* | KLF13 | MAFF | STO | CDF3 |
| 18 | RTT107 | WTM2 | NR0B2 | GM6710 | PIL6 | ABF1 |
| 19 | ESC2 | CSE2 | ZFP36l1 | EN2 | CCA1* | TOC1* |
| 20 | ASF1 | GAT1 | ATF5 | ZBTB7C | EPR1 | EPR1 |
| 21 | WTM2 | OTU1 | SREBF1 | MESP1 | LZF1 | AT1G70000 |
| 22 | HAP5 | HCM1* | STAT5B | NR1D2* | BZS1 | STO |
| 23 | HPA2 | PHD1 | GTF2IRD1 | ZFP987 | ASG4 | BBX16 |
| 24 | EDS1 | MAL13 | ZBTB21 | GM14401 | CDF3 | ATCTH |
| 25 | RTG2 | RGT2 | SOX9 | GM14305 | CO | LUX* |
| Recall | 35.3% | 47.1% | 66.7% | 35.7% | 45.5% | 54.5% |

LL_LDHC: Constant light and temperature; LDHC: 24 hour cycling light and temperature; MA: Microarray; RNA: RNAseq

* Core transcription factors in Dataset S3.

**Table S3. The top 25 highest PerReg-ranked genes among all transcription factors.**

| Rank | *S. cerevisiae* | | *M. Musculus* | | *A. thaliana* | |
|---|---|---|---|---|---|---|
| | MA | RNA | MA | RNA | LDHC | LL_LDHC |
| 1 | *RME1* | ***YOX1**** | ***NR1D1**** | *CDX* | *RVE1* | *MYBL2* |
| 2 | ***ASH1**** | ***TOS4**** | ***NPAS2**** | ***NPAS2**** | ***LHY**** | ***CCA1**** |
| 3 | ***SWI5**** | *RME1* | ***DBP**** | *GM14444* | ***CCA1**** | ***LHY**** |
| 4 | *HST3* | ***ASH1**** | ***ARNTL**** | *FOXO6* | *COL2* | *COL2* |
| 5 | ***ACE2**** | ***SWI5**** | *EGR1* | ***NR1D1**** | *STH* | ***PRR7**** |
| 6 | ***SWI4**** | *KAR4* | ***NR1D2**** | *POU4F1* | *MYBL2* | ***PRR5**** |
| 7 | ***YOX1**** | *HST3* | *MYC* | *GM20422* | *AT1G26790* | *PIF4* |
| 8 | *HST4* | *ZNF1* | ***NFIL3**** | *DMRTA2* | ***PRR7**** | *STH* |
| 9 | *SIP4* | *RTT107* | ***BHLHE41**** | *NPAS3* | *HYH* | *COL5* |
| 10 | *ASF1* | *MGA1* | ***TEF**** | *EGR1* | ***PRR5**** | *RVE1* |
| 11 | ***TOS4**** | ***SWI4**** | ***RORC**** | ***DBP**** | *BBX16* | *CDF3* |
| 12 | *RTT107* | *SIP4* | *ZBTB16* | *CREB5* | *LZF1* | *LZF1* |
| 13 | ***STB1**** | ***ACE2**** | *PPARD* | *TBX1* | *BBX18* | ***LUX**** |
| 14 | *EDS1* | *PHD1* | *MAFB* | *EGR3* | *CDF2* | *CDF1* |
| 15 | *GAT1* | *MAL33* | *KLF10* | *MESP1* | *EPR1* | *COL1* |
| 16 | *HAP4* | *HST4* | *NR0B2* | *GM6710* | *CDF1* | *ATCTH* |
| 17 | *HPA2* | *TEC1* | *RFXANK* | ***ARNTL**** | *COL1* | *CDF2* |
| 18 | *GAL3* | *RLF2* | *BCl6* | *EN2* | *BBX19* | *BBX18* |
| 19 | *WTM1* | *ASF1* | *ZFP931* | *ZBTB7C* | ***LUX**** | *ABF1* |
| 20 | *RLF2* | ***FKH1**** | *NR3C2* | *GM14401* | *PIF4* | *COL9* |
| 21 | *NDT80* | *MSS11* | ***CLOCK**** | *GM14305* | ***PRR9**** | *EPR1* |
| 22 | *XBP1* | *MTH1* | *ONECUT1* | *BHLHE22* | *CDF3* | *AT1G26790* |
| 23 | *TEC1* | *RPI1* | *KLF9* | *PPARD* | *RVE2* | *BBX16* |
| 24 | *MSN4* | ***STB1**** | *ZBTB7C* | *OVOl2* | ***RVE8**** | *AT5G44260* |
| 25 | ***HCM1**** | ***HCM1**** | *FOXQ1* | *FOXP3* | *TGA3* | *HYH* |
| Recall | 47.1% | 52.9% | 66.7% | 28.6% | 63.6% | 45.5% |

LL_LDHC: Constant light and temperature; LDHC: 24 hour cycling light and temperature; MA: Microarray; RNA: RNAseq
***** Core transcription factors in Dataset S3.

#### 3 Figures

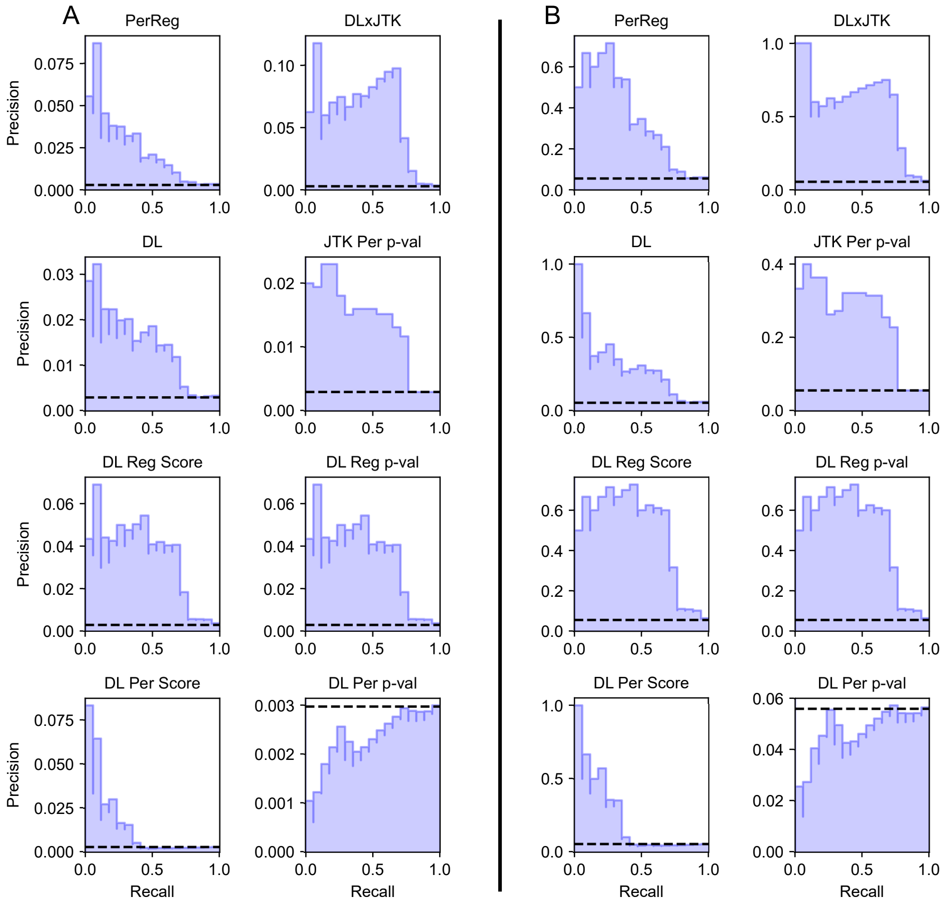


**Fig S1. Precision-recall curves of classifiers identifying core from non-core TFs among all genes (A) and among only TFs (B) in *S. cerevisiae* microarray dataset.** Between changes in recall, $R_{n-1} < R_n$, precision is plotted as a constant equal to the precision $P_n$ at the minimum decision-threshold rank giving a recall of $R_n$, so that the area under the curve is equal to average precision. The horizontal dashed line indicates the baseline average precision of a random classifier.

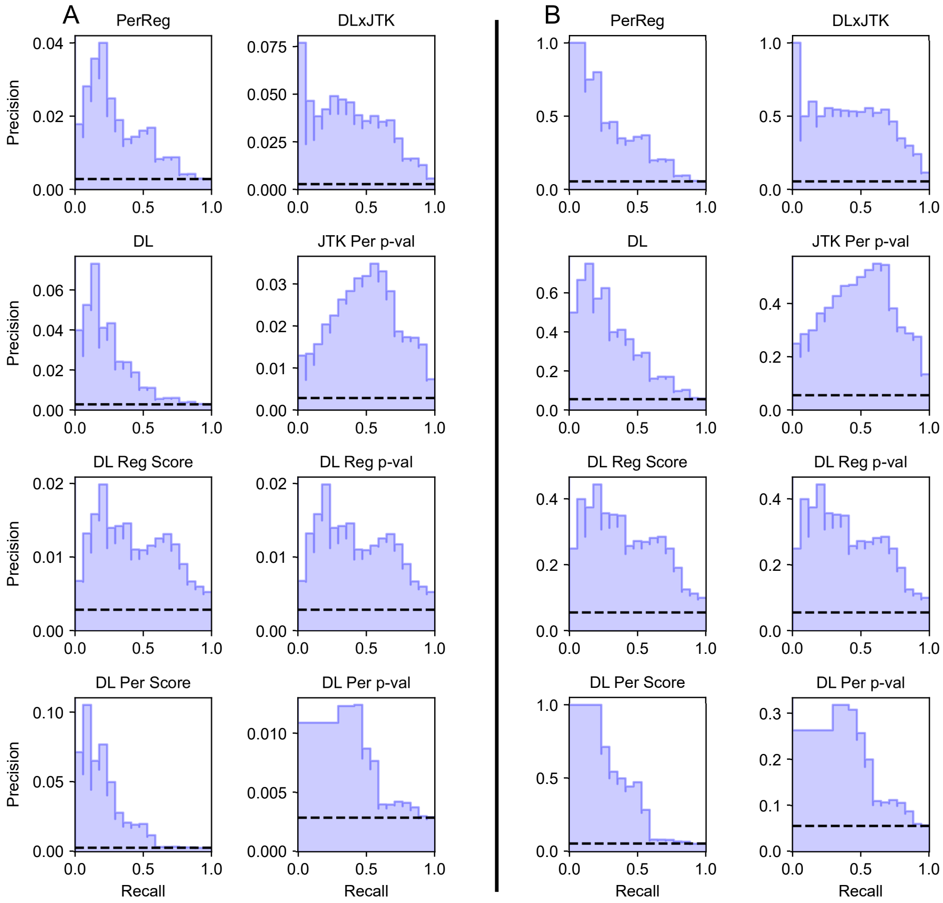


**Fig S2. Precision-recall curves of classifiers identifying core from non-core TFs among all genes (A) and among only TFs (B) in *S. cerevisiae* RNAseq dataset.** Between changes in recall, $R_{n-1} < R_n$, precision is plotted as a constant equal to the precision $P_n$ at the minimum decision-threshold rank giving a recall of $R_n$, so that the area under the curve is equal to average precision. The horizontal dashed line indicates the baseline average precision of a random classifier.

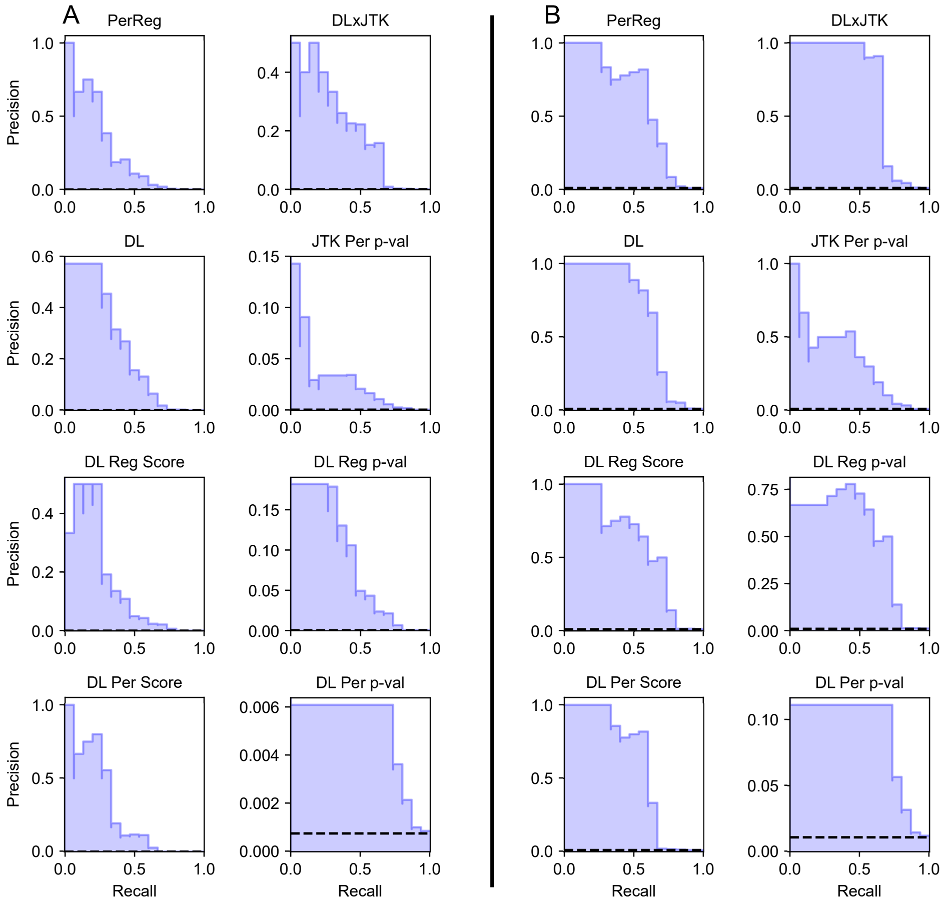


**Fig S3. Precision-recall curves of classifiers identifying core from non-core TFs among all genes (A) and among only TFs (B) in *M. Musculus* microarray dataset.** Between changes in recall, $R_{n-1} < R_n$, precision is plotted as a constant equal to the precision $P_n$ at the minimum decision-threshold rank giving a recall of $R_n$, so that the area under the curve is equal to average precision. The horizontal dashed line indicates the baseline average precision of a random classifier.

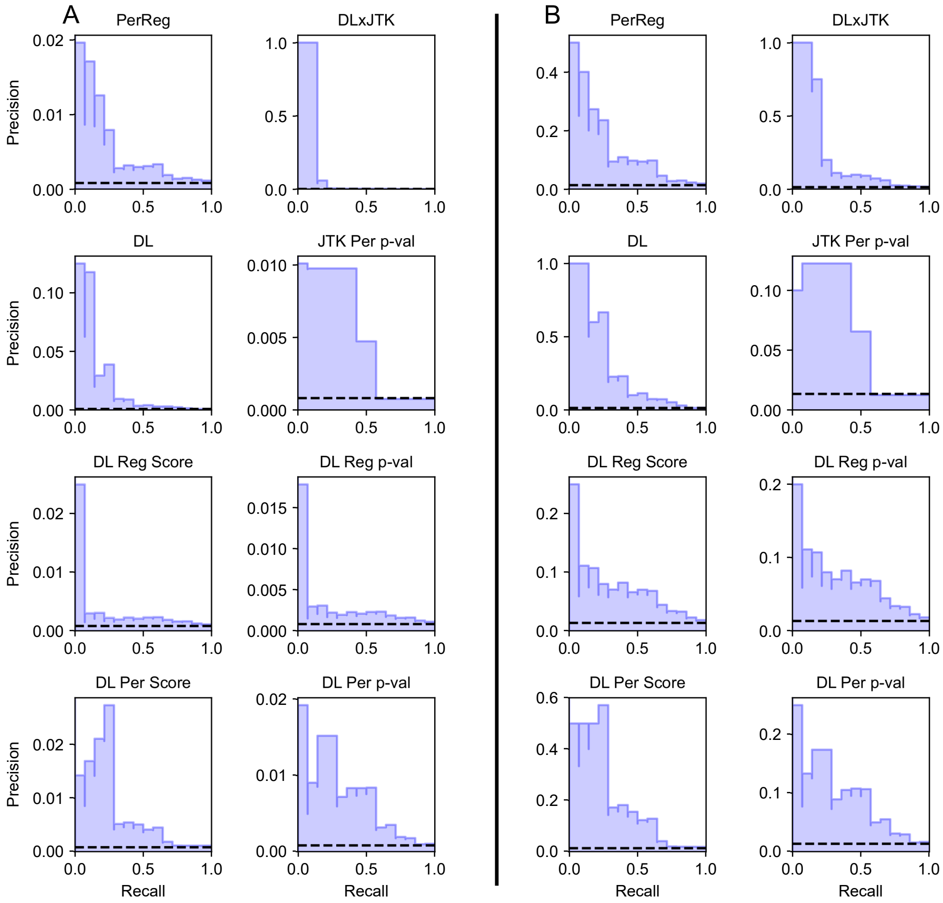


**Fig S4. Precision-recall curves of classifiers identifying core from non-core TFs among all genes (A) and among only TFs (B) in *M. Musculus* RNAseq dataset.** Between changes in recall, $R_{n-1} < R_n$, precision is plotted as a constant equal to the precision $P_n$ at the minimum decision-threshold rank giving a recall of $R_n$, so that the area under the curve is equal to average precision. The horizontal dashed line indicates the baseline average precision of a random classifier.

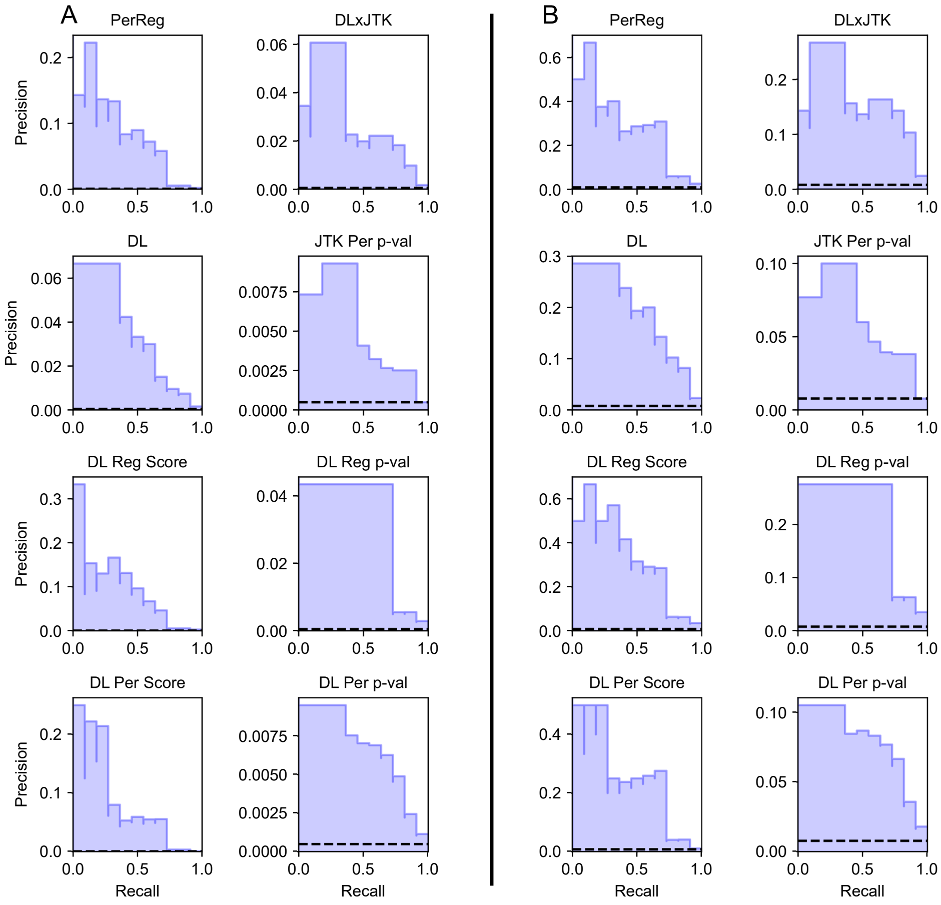


**Fig S5. Precision-recall curves of classifiers identifying core from non-core TFs among all genes (A) and among only TFs (B) in *A. thaliana* microarray LDHC dataset.** Between changes in recall, $R_{n-1} < R_n$, precision is plotted as a constant equal to the precision $P_n$ at the minimum decision-threshold rank giving a recall of $R_n$, so that the area under the curve is equal to average precision. The horizontal dashed line indicates the baseline average precision of a random classifier.

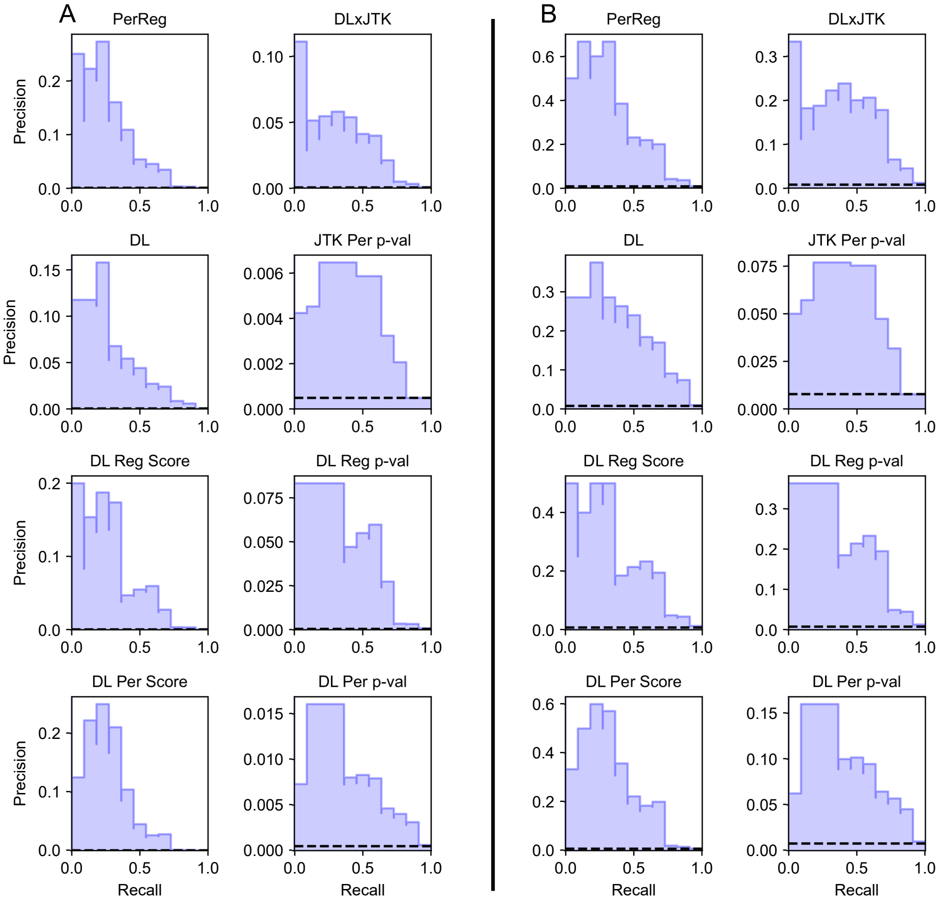


**Fig S6. Precision-recall curves of classifiers identifying core from non-core TFs among all genes (A) and among only TFs (B) in *A. thaliana* microarray LL_LDHC dataset.** Between changes in recall, $R_{n-1} < R_n$, precision is plotted as a constant equal to the precision $P_n$ at the minimum decision-threshold rank giving a recall of $R_n$, so that the area under the curve is equal to average precision. The horizontal dashed line indicates the baseline average precision of a random classifier.

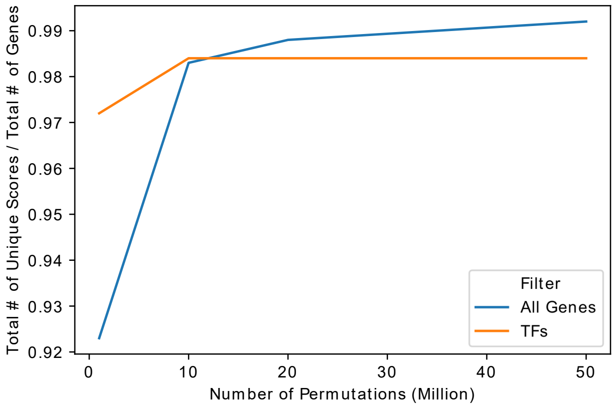


**Fig S7. The number of permutations for computing DL Reg $p$-val vs. the number of unique DL Reg $p$-values.** The number of unique values of DL Reg $p$-val per total number of genes before filtering for transcription factors (blue line) and after filtering for transcription factors (orange line) for the *A. thaliana* LDHC dataset as a function of the number of permutations chosen for the DL algorithm.

### References


1. de Lichtenberg U, Jensen LJ, Fausbøll A, Jensen TS, Bork P, Brunak S. Comparison of computational methods for the identification of cell cycle-regulated genes. Bioinformatics. 2005;21(7):1164–1171. doi:10.1093/bioinformatics/bti093.

2. Motta F, Deckard A, Moseley RC. pyDL; 2019. `https://gitlab.com/biochron/pydl`.

3. Hughes ME, Hogenesch JB, Kornacker K. JTK_CYCLE: An Efficient Nonparametric Algorithm for Detecting Rhythmic Components in Genome-Scale Data Sets. Journal of Biological Rhythms. 2010;25(5):372–380. doi:10.1177/0748730410379711.

4. Kendall MG. A NEW MEASURE OF RANK CORRELATION. Biometrika. 1938;30(1-2):81–93. doi:10.1093/biomet/30.1-2.81.

5. Harding EF. An Efficient, Minimal-Storage Procedure for Calculating the Mann-Whitney U, Generalized U and Similar Distributions. Journal of the Royal Statistical Society Series C (Applied Statistics). 1984;33(1):1–6.

6. Hutchison A. pyJTK: Python implementation of the JTK_CYCLE statistical test; 2013. `https://github.com/alanlhutchison/pyJTK`.

7. Hutchinson A, Motta F, Deckard A. pyJTK; 2019. `https://gitlab.com/biochron/pyjtk`.
